# iRhom2 regulates ERBB signalling to promote KRAS-driven oncogenesis

**DOI:** 10.1101/2021.08.06.455383

**Authors:** Boris Sieber, Fangfang Lu, Stephen M. Stribbling, Adam G. Grieve, Anderson J. Ryan, Matthew Freeman

## Abstract

Dysregulation of the ERBB/EGFR signalling pathway causes multiple types of cancer (1, 2). Accordingly, ADAM17, the primary shedding enzyme that releases and activates ERBB ligands, is tightly regulated. It has recently become clear that iRhoms, inactive members of the rhomboid-like superfamily, are regulatory cofactors for ADAM17 (3, 4). Here we show that oncogenic KRAS mutants target the cytoplasmic domain of iRhom2 to induce ADAM17-dependent shedding and the release of ERBB ligands. Activation of ERK1/2 by oncogenic KRAS induces the phosphorylation of iRhom2, recruitment of the phospho-binding 14-3-3 proteins, and consequent ADAM17-dependent shedding of ERBB ligands. In addition, cancer-associated mutations in iRhom2 act as sensitisers in this pathway by further increasing KRAS-induced shedding of ERBB ligands. This mechanism is conserved in lung cancer cells, where iRhom activity is required for tumour xenograft growth. In this context, the activity of oncogenic KRAS is modulated by the iRhom2-dependent release of ERBB ligands, thus placing iRhom2 as a central component of a positive feedback loop in lung cancer cells. Overall, the cytoplasmic domain of iRhom2 is a critical component of KRAS-induced oncogenesis of lung cancer cells. Both ADAM17 and iRhom2 have also been implicated in a wide range of other cancers (5–10), so the mechanism we have revealed may also have wider oncogenic significance.

## Introduction

The ERBB/EGFR signalling pathway is dysregulated in numerous cancers, especially of the lung, breast and ovary (1, 2). In addition to oncogenic receptor mutations, tumorigenesis can be driven by excess ERBB ligand production (11). ERBB family ligands are mostly synthesised as type I transmembrane domain proteins, and become active upon proteolytic cleavage and release (shedding) from the plasma membrane. Thus, shedding of ERBB ligands is a primary regulator of signalling that controls pathogenesis as well as cell proliferation, survival and differentiation.

This mode of regulation puts into the spotlight the enzymes responsible for shedding ligands. The metalloprotease ADAM17 is the most widespread sheddase of ERBB ligands, as well as controlling the release of many other growth factors, cytokines and other cell surface proteins (12). Consistent with its potency, an intricate regulatory mechanism exists to control ADAM17, centred on iRhom1 and iRhom2, which are rhomboid-like proteins that act as ADAM17 cofactors (4). For example, iRhom2 is required for the maturation and subsequent activation at the plasma membrane of ADAM17 to catalyse the shedding of TNFα, the primary inflammatory cytokine. This plasma membrane activation of ADAM17 can be triggered by ERK1/2-dependent phosphorylation of the cytoplasmic domain of iRhom2 (13, 14).

Several lines of evidence have implicated iRhoms and ADAM17 in tumorigenesis, especially in lung, breast, cervical, oesophageal and colorectal cancers. iRhom2 and ADAM17 levels increase during cancer progression and correlate with lower survival rates (5–7, 15–18). The most direct link between the iRhoms and cancer are mutations in the cytoplasmic domain of iRhom2, which cause a rare familial syndrome, tylosis with oesophageal cancer (TOC), characterised by a very high lifetime risk of developing oesophageal cancer (8, 19–22). Increased activity of ADAM17 has been observed for *iRhom2^TOC^* mutations (23, 24) but, despite this strong genetic link, the precise mechanistic role of iRhoms in oncogenic signalling has been poorly explored.

ADAM17 is better characterised than iRhoms with respect to cancer, although until recently it too has not been the subject of the intense focus commensurate with its regulatory importance. For instance, oncogenic SRC triggers the ADAM17-dependent release of the ERBB ligand TGFα (25). It has also become clear that ADAM17 is important in cancers mediated by mutations in *KRAS*, which are the most frequent oncogenic mutations in human cancers, particularly in lung, colorectal and pancreatic tumours (26). Although oncogenic KRAS has long been considered to be constitutively active, and thus independent of upstream signals, a requirement for ADAM17 and ERBB1/EGFR in KRAS-induced pancreatic cancer has challenged this idea (27, 28). Indeed, ERBB signalling has now been shown to contribute to lung tumorigenesis by supporting activation of oncogenic KRAS (29, 30). In this context, it is also significant that KRAS-driven tumours express higher levels of ERBB ligands, in particular amphiregulin and TGFα (27, 29). However, as described above, ERBB ligands must be proteolytically shed to be active, and the regulation of shedding in cancer has been largely unknown. A recent advance has been the demonstration of a requirement for ADAM17 in KRAS-induced lung tumorigenesis (31). Using NSCLC and patient-derived xenografts, as well as the *Kras^G12D^* mouse model, Saad et al. showed that depletion of ADAM17, or inhibition of its activity, suppressed lung tumour growth. They also found that oncogenic KRAS leads to increased p38 activity, which induces the phosphorylation of ADAM17, a marker of its activity (3, 12), as well as causing upregulated shedding of the ADAM17 substrate IL-6R.

Here, we report that that iRhoms are essential for the oncogenic release of ERBB ligands by KRAS-G12 mutants. Specifically, KRAS-induced shedding of ERBB ligands is triggered by the phosphorylation of the cytoplasmic domain of iRhom2, which allows the recruitment of the phospho-binding proteins 14-3-3. Human cancer-associated mutations in the cytoplasmic domain of iRhom2 are sufficient to amplify this pathway, thus further establishing iRhom2 as an important component of oncogenic signalling. The pathological significance of this pathway was validated upon oncogenic KRAS expression in HEK239T cells and in non-small-cell lung carcinoma (NSCLC) cell line A549 harbouring an endogenous oncogenic *KRAS^G12S^* mutation. Furthermore, loss of iRhom activity completely suppressed KRAS-driven tumour xenograft growth, demonstrating the requirement of iRhoms in a widely used model of lung cancer. Finally, we report that the cytoplasmic domain of iRhom2 is a hub for an ERBB-dependent positive feedback loop that maintains KRAS activity in lung cancer cells. Overall, our results demonstrate that iRhom2 plays a central role in oncogenic KRAS-induced signalling.

## Results

### iRhoms are required for KRAS-driven shedding of ERBB ligands by ADAM17

Oncogenic KRAS induces the activation of ADAM17 (27, 31, 32), so we questioned whether iRhoms play a role in this process. First, to establish the effect of oncogenic KRAS in HEK293T cells, we expressed KRAS^G12V^. As expected, we observed a significant increase in the release of the ADAM17 substrate TGFα (Fig. 1A) especially compared to the effect of KRAS^S17N^ (Fig. 1A), a mutant with reduced GTPase activity (33, 34). Using the inhibitors GI254023X and GW280264X, which respectively inhibit ADAM10, or ADAM10 and ADAM17 (35), we confirmed that TGFα shedding by oncogenic KRAS was dependent on ADAM17 (Fig. 1B), which agrees with the reported ability of oncogenic KRAS to increase ADAM17-dependent shedding (27, 31, 32).

**Figure 1.**
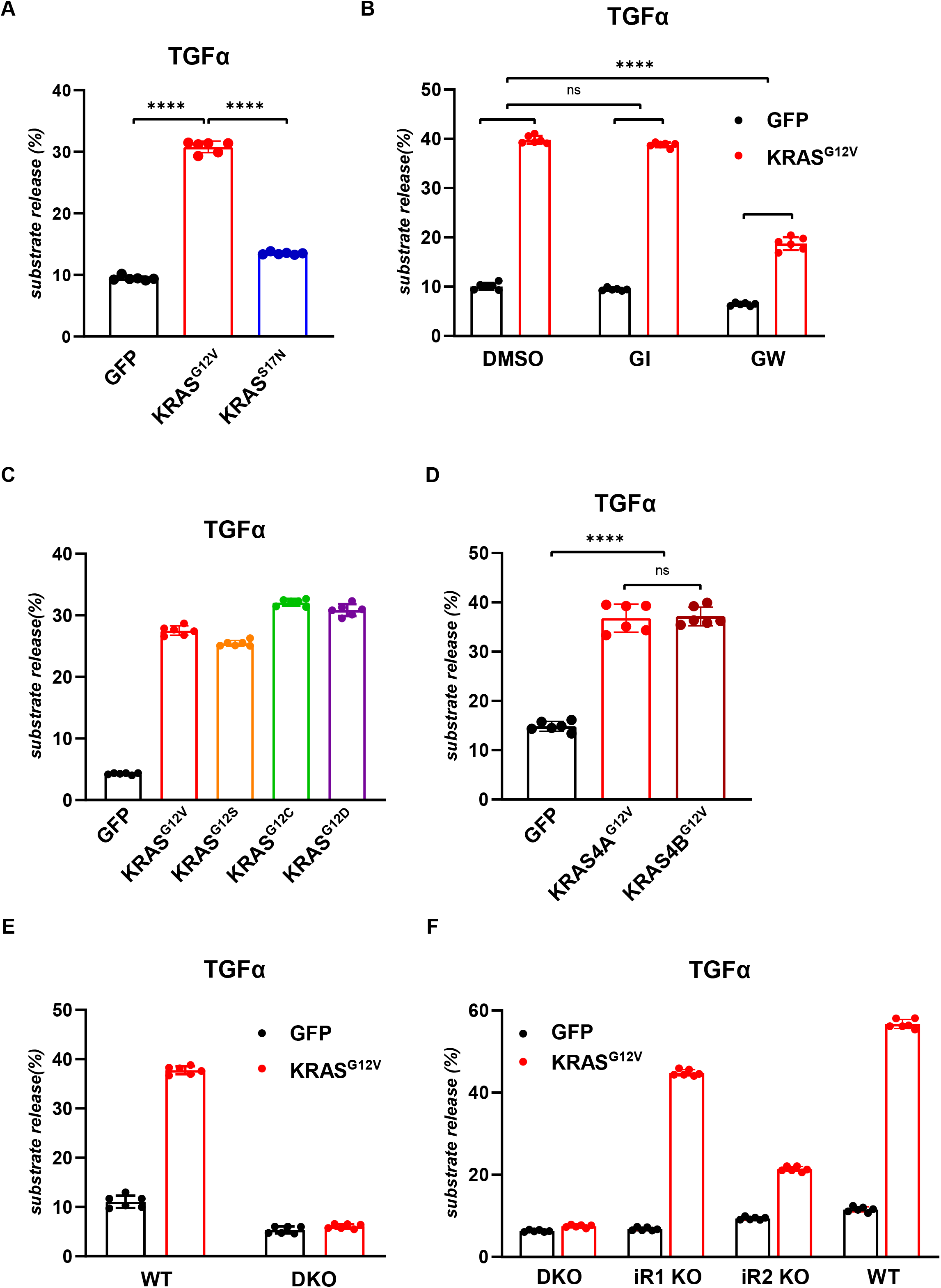
iRhoms are required for KRAS-driven shedding of the ERBB ligands by ADAM17. **A-D.** HEK293T cells were co-transfected with alkaline-phosphatase (AP)-tagged TGFα and GFP or GFP-tagged KRAS constructs. Unless specified otherwise, constructs of KRAS4A were used in all experiments. Overnight medium collection was performed in presence of 0.5 μM ADAM10 inhibitor (GI), 0.5 μM ADAM10/ADAM17 inhibitor (GW) or with DMSO. **E-F.** Wild-type, iRhom1 KO, iRhom2 KO, or iRhom1/2 double knockout (DKO) HEK293T cells were transiently co-transfected with AP-tagged TGFα and GFP or GFP-tagged KRAS-G12V, followed by overnight medium collection. Substrate release is the level of alkaline phosphatase in the medium divided by the total alkaline phosphatase level. Data are from six biological replicates. Error bars represent standard deviations and statistical tests were performed using two-tailed student t-test. ns = p value>0.05, **** = p value<0.0001.

Although KRAS^G12V^ is one of the most well studied oncogenic forms of KRAS (36), several other *KRAS^G12X^* mutations are found in human cancers (37, 38). We found that KRAS^G12S^, KRAS^G12C^ and KRAS^G12D^ all caused elevated TGFα release (Fig. 1C). We also demonstrated that both isoforms of KRAS, 4A and 4B, induce shedding of TGFα (Fig. 1D). Overall, these results demonstrate the shared ability of KRAS oncogenic mutants to trigger growth factor release.

Having shown that oncogenic mutations in KRAS induce ADAM17-dependent shedding of TGFα, we next asked whether iRhoms are required for this activity. We found that KRAS-induced shedding of TGFα was completely blocked in HEK293T double-knockout (DKO) cells mutant for both iRhom1 and iRhom2 (Fig. 1E, S1A). In single knockout lines, loss of iRhom1 had little effect, whereas iRhom2 KO showed a strong reduction in TGFα shedding (Fig. 1F), thereby demonstrating that iRhom2 is the primary mediator of KRAS-induced ADAM17-dependent shedding of TGFα.

### KRAS-induced shedding depends on phosphorylation of the cytoplasmic domain of iRhom2 by the Raf/MEK/ERK pathway

To determine whether, as in inflammatory signalling (13, 14), iRhom2 phosphorylation participates in oncogenic ADAM17 signalling, we used a mutant version of iRhom2 (iRhom2^site1-3^) in which the three primary phosphorylation sites are changed to alanine (14). We found that without iRhom2 phosphorylation at these three main sites, shedding of TGFα was significantly reduced (Fig. 2A). Importantly, this phosphorylation-deficient form of iRhom2 supported ADAM17 maturation as efficiently as iRhom2^WT^ (Fig. S2A), which aligns with our previous findings that iRhom2 phosphorylation is not needed for ADAM17 maturation (14). These results reveal the role of iRhom2 phosphorylation in oncogenic signalling by KRAS.

**Figure 2.**
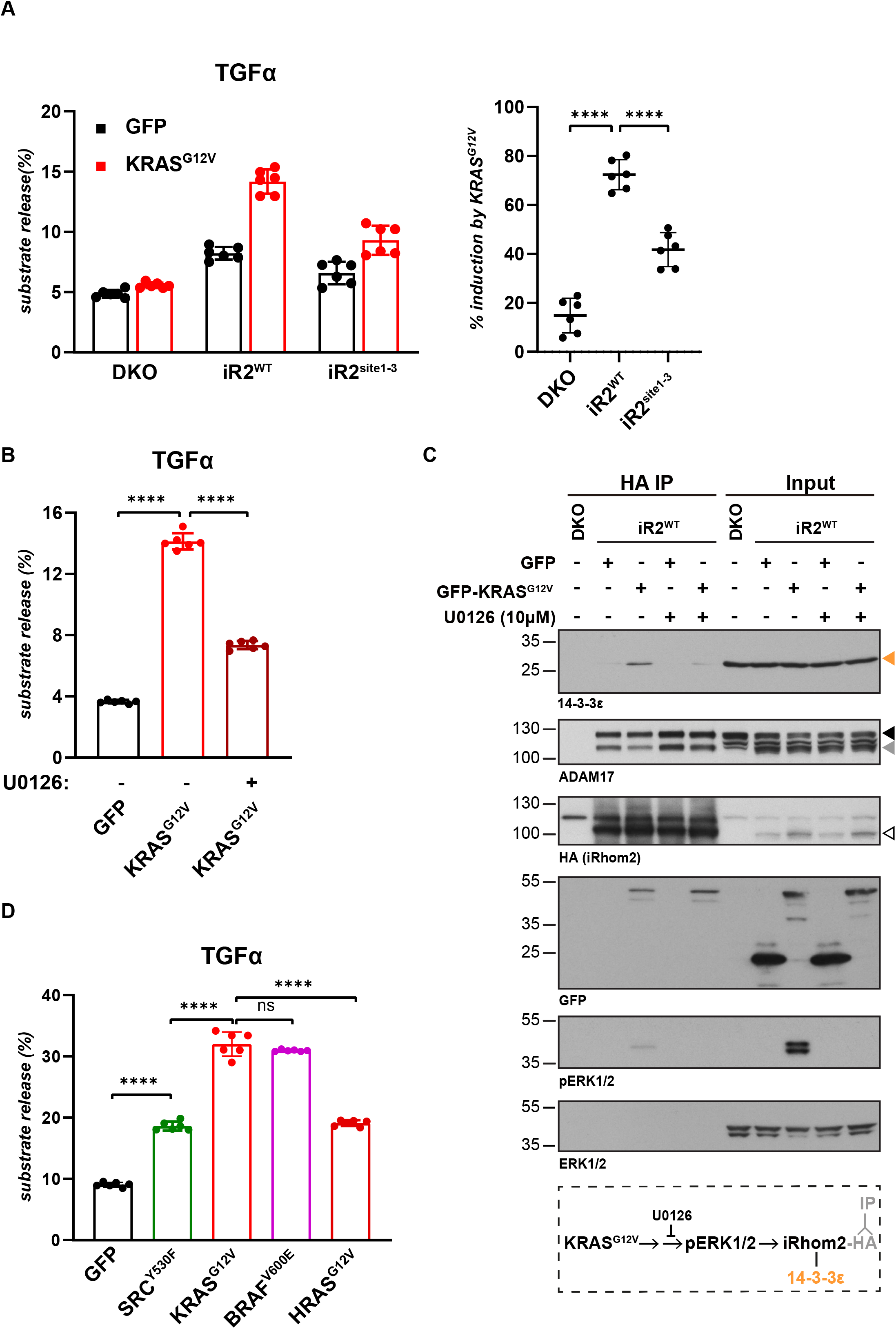
KRAS-induced shedding depends on the phosphorylation of the cytoplasmic domain of iRhom2 by the Raf/MEK/ERK pathway. **A.** iRhom1/2 DKO HEK293T cells reconstituted with iRhom2^WT^ and iRhom2 lacking the three primary phosphorylation sites (iRhom2^site1-3^) were co-transfected with GFP or GFP-tagged KRAS^G12V^ and alkaline-phosphatase (AP)-tagged TGFα, and medium was collected overnight. The right panel shows the percentage of AP release induced by KRAS^G12V^ in the indicated cell lines. Data are from six biological replicates. **B.** iRhom1/2 DKO HEK293T reconstituted with iRhom2^WT^ and co-transfected with GFP or GFP-tagged KRAS^G12V^ and AP-TGFα were treated with 10 μM U0126 during three hours of medium collection. Data are from six biological replicates. **C.** HA-based immunoprecipitates and lysates from iRhom1/2 DKO HEK293T reconstituted with HA-iRhom2^WT^ and transfected with GFP or GFP-tagged KRAS^G12V^ were immunoblotted for 14-3-3ε, ADAM17, HA and GFP. To assess the contribution of Raf/MEK/ERK cascade, cells were treated with 10 μM U0126 for two hours and blotted for phosphorylated ERK1/2 (pERK1/2). Orange arrowhead indicates 14-3-3ε, black and grey arrowheads indicate immature proADAM17 and mature ADAM17 respectively, white arrowhead indicates HA-tagged iRhom2^WT^. The experiment was repeated three times. A schematic of the rationale of the experiment is shown below the immunoblot. **D.** HEK293T cells were transiently co-transfected with AP-tagged TGFα and GFP or GFP-tagged ERK-activating oncogenes SRC^Y530F^, KRAS^G12V^, BRAF^V600E^, HRAS^G12V^, followed by overnight medium collection. Data are from six biological replicates. Substrate release is the level of released alkaline phosphatase in the medium divided by the total alkaline phosphatase level. Error bars represent standard deviations and statistical tests were performed using two-tailed student t-test. ns = p value>0.05, **** = p value<0.0001.

In inflammatory signalling, iRhom2 phosphorylation is MAP kinase dependent (13, 14); it is also well established that oncogenic KRAS mutations act through the Raf/MEK/ERK MAP kinase pathway (36, 39). We therefore asked whether the RAS/MAPK cascade also participates in iRhom2-dependent oncogenic signalling. TGFα shedding induced by KRAS^G12V^ was strongly inhibited by treating the cells with U1026 (Fig. 2B), a specific inhibitor of MEK1/2 (40), the kinases upstream of ERK1/2. We also found that oncogenic KRAS triggers the recruitment of 14-3-3 epsilon to iRhom2 and that, consistent with 14-3-3 proteins binding to phosphorylated residues (41), this recruitment was inhibited by treatment with U1026 (Fig. 2C). 14-3-3 recruitment to iRhom2 was associated with decreased binding between iRhom2 and ADAM17 (Fig. 2C). Although we have not investigated this phenomenon further it agrees with our previous work on inflammatory signalling (14) and suggests that the activation of ADAM17 by phosphorylated iRhom2 depends on an altered interaction between them. Since the recruitment of 14-3-3 to iRhom2 is sufficient for ADAM17 activation (13, 14), these results demonstrate that KRAS-induced shedding of ERBB ligands is mediated by ERK1/2-dependent phosphorylation of iRhom2.

ERK1/2 activation is not only induced by oncogenic KRAS but also by several other oncogenes (42–44), so we asked whether these other ERK1/2 activating oncogenes can similarly drive ADAM17 activity. HRAS^G12V^, BRAF^V600E^ and SRC^Y530F^, all of which activated ERK1/2 (Fig. S2B), also induced elevated release of TGFα from HEK293T cells (Fig. 2D). This result is consistent with an increase in TGFα release by oncogenic SRC (25). ERK-activating oncogenes KRAS^G12V^ and BRAF^V600E^ also triggered the release of amphiregulin (Fig. S2C), another ADAM17-dependent ERBB ligand with a well-established role in oncogenesis (45, 46). This contrasted with no increase of pERK levels (Fig. S2B) (47, 48) and little effect on amphiregulin release (Fig. S2C) caused by the oncogene AKT^E17K^. These results suggest that the ability of ERK1/2-activating oncogenes to trigger the release of ERBB ligands depends on a common mechanism driven by phosphorylated iRhom2.

### Cancer-associated mutations in iRhom2 potentiate KRAS-induced shedding of ERBB ligands

Our data demonstrate that iRhom2 phosphorylation participates in oncogenic signalling. The strongest and most direct evidence for the involvement of iRhom2 in human cancer is in the case of a rare inherited syndrome called tylosis with oesophageal cancer (TOC), which is caused by mutations in a small and highly conserved region within the cytoplasmic N-terminal domain of iRhom2 (Fig. 3A) (8). TOC is characterised by hyperkeratosis, oesophageal cancer, and at least in the case of one of the familial mutations, iRhom2^D188N^, by a susceptibility to other cancers (22). We therefore investigated whether the tylotic mutations affect oncogenic signalling through ADAM17. Replacing wild-type iRhom2 with tylotic iRhom2^D188N^ caused a strong enhancement of KRAS-induced shedding of the ADAM17 substrate and ERBB ligand amphiregulin (Fig. 3B). The shedding of EGF, which is triggered by ADAM10 rather than ADAM17 (49), is not affected by iRhom2^D188N^ (Fig. 3B), demonstrating the specificity of the oncogenic iRhom2 mutation for ADAM17. Strikingly, all analysed TOC mutations, including when combined, amplified KRAS-induced amphiregulin release (Fig. 3C); none affected EGF shedding (Fig. S3B). Furthermore, none of the tylotic mutations altered ADAM17 maturation (Fig. S3A), consistent with our previous conclusion that the cytoplasmic tail of iRhom2 does not participate in the earlier iRhom2 function of promoting ER to Golgi trafficking of ADAM17 (14). We conclude that TOC mutations are sufficient to potentiate KRAS-induced shedding of ADAM17 substrates in HEK293T cells, thereby establishing the direct effect of mutations in N-terminus of iRhom2 in oncogene-driven signalling.

**Figure 3.**
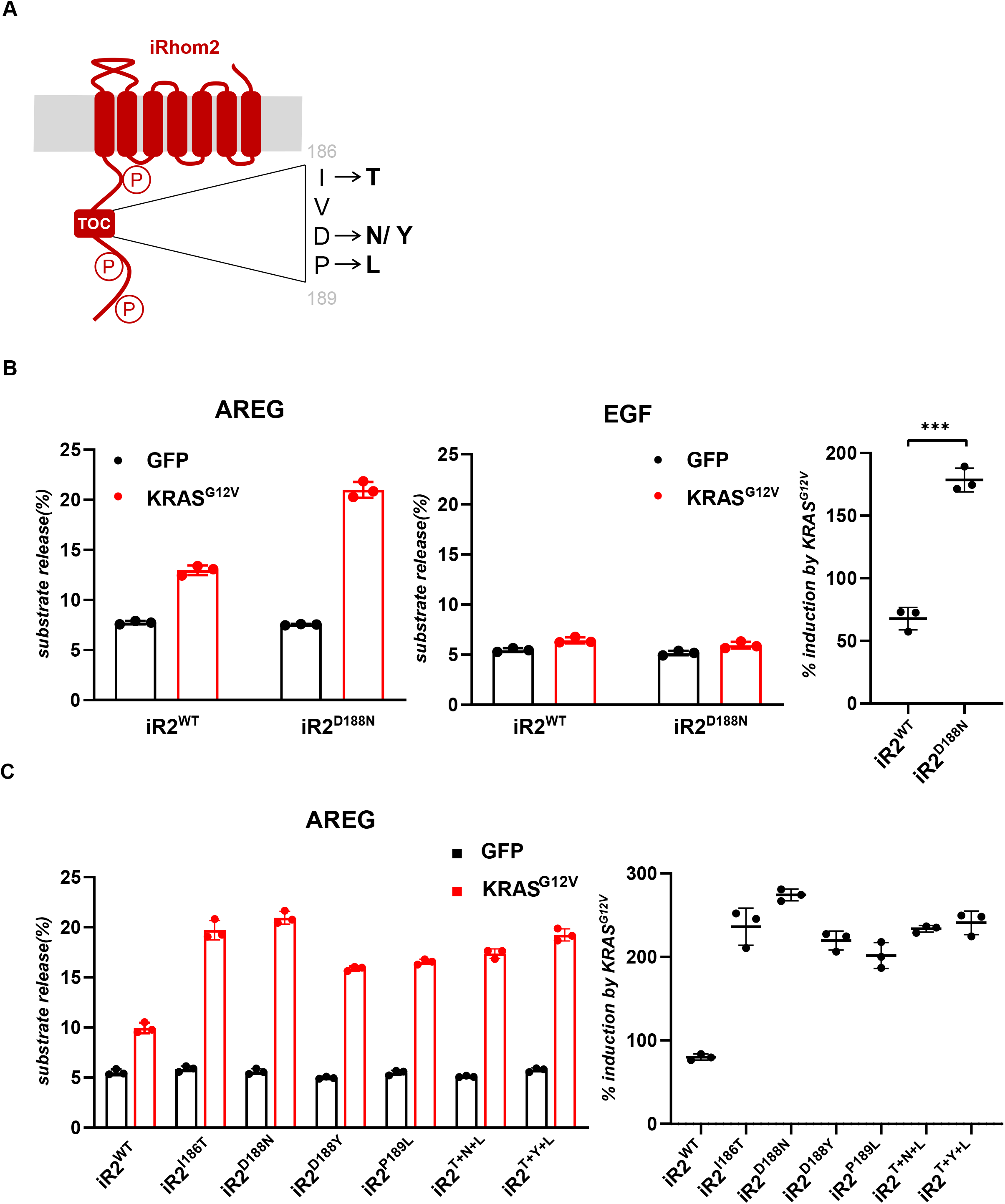
Cancer-associated mutations in iRhom2 potentiate KRAS-induced shedding of ERBB ligands. **A.** Schematic of the N-terminal domain of iRhom2 with the three main sites of phosphorylation and the conserved region that harbours mutations causing tylosis with oesophageal cancer (TOC). The four analysed TOC mutations are shown in the insert. **B-C.** iRhom1/2 DKO HEK293T cells were reconstituted with iRhom2^WT^ or with an iRhom2 variant harbouring one of the TOC mutations or the three mutations combined: T+Y+L (I186T, D188Y, P189L) or T+N+L (I186T, D188N, P189L). Upon co-transfection with GFP or GFP-tagged KRAS^G12V^, and alkaline-phosphatase (AP)-tagged AREG or EGF, overnight collection of medium was performed in biological triplicates. Substrate release is the level of released alkaline phosphatase in the medium divided by the total alkaline phosphatase level. The far right panels show the percentage of AP-AREG release induced by KRAS^G12V^ in the indicated cell lines. Error bars represent standard deviations and statistical tests were performed using two-tailed student t-test. *** = p value<0.001.

### iRhoms are required for KRAS-driven tumorigenesis

Increased activation of ERBB1/EGFR as well as of the other ERBB receptors have widespread involvement in cancers (1, 50, 51) including, it has recently been established, in KRAS-induced lung tumorigenesis (29, 30). We therefore addressed the potential role of iRhoms in A549 cells, a widely used human lung adenocarcinoma cell model. These cells were selected because they are homozygous for *KRAS^G12S^*, one of the mutations that we have shown to drive TGFα release (Fig 1C). Using CRISPR/Cas9, we knocked out both *iRhom1* and *iRhom2* in A549 cells to create a A549-DKO cell line, lacking all iRhom activity (Fig. S4A). Consistent with our data from HEK293T cells, loss of iRhoms abolished all shedding of the endogenous ERBB ligand amphiregulin (Fig. 4A), demonstrating that iRhoms promote growth factor signalling in a lung cancer cell line. In support of this conclusion, DKO cells also showed a decrease in cell proliferation (Fig. S4B).

**Figure 4.**
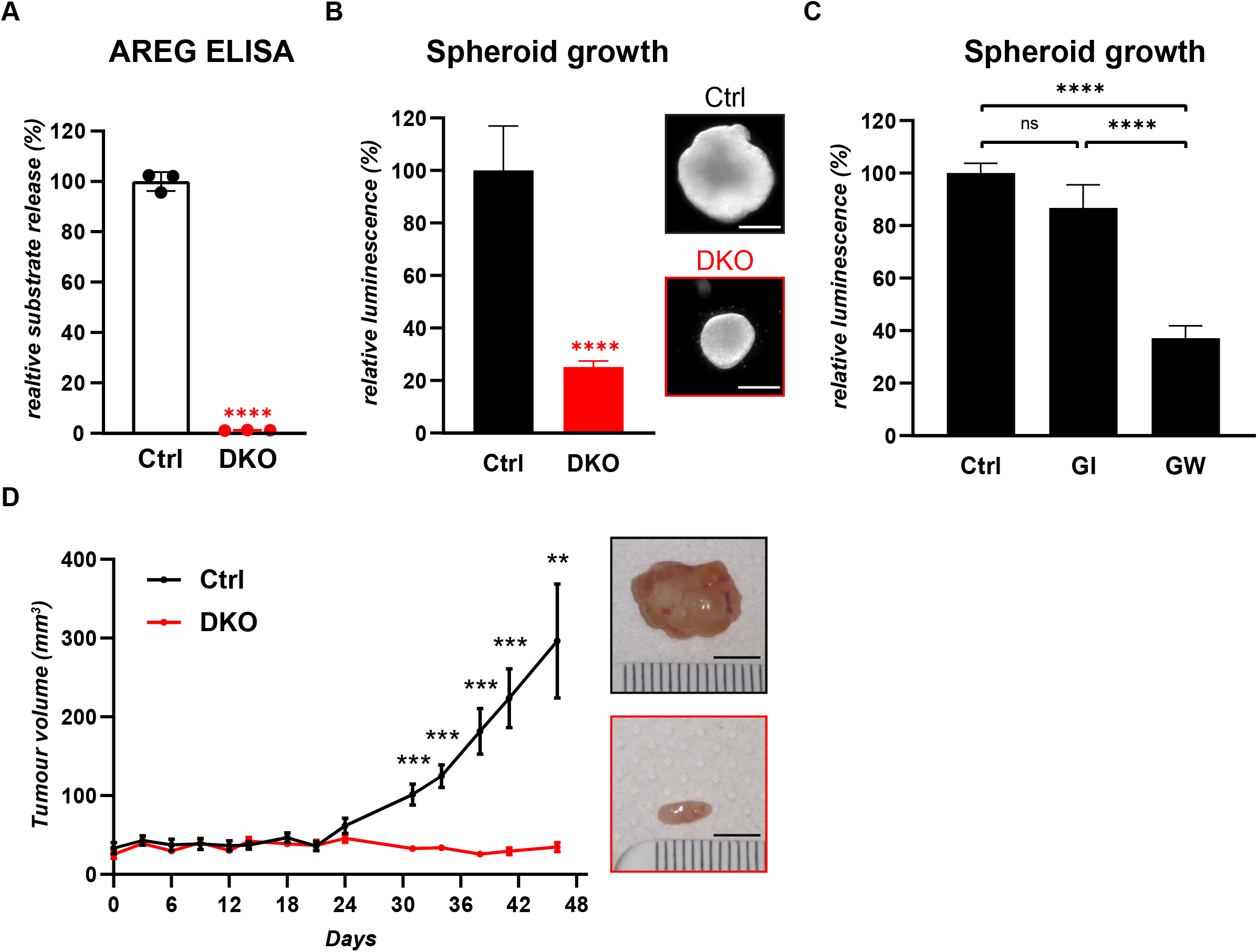
iRhoms are required for KRAS-driven tumorigenesis. **A.** Release of endogenous amphiregulin (AREG) from control (Ctrl) and iRhom1/2 DKO lung cancer cells A549 was measured after overnight collection in biological triplicates. AREG concentration determined by ELISA was normalised to the total protein concentration in A549 cells, and the average level of A549 Ctrl was defined as the reference (100%). Unless specified otherwise, ELISA experiments are similarly normalised in all experiments. Error bars represent standard deviations and statistical tests were performed using two-tailed student t-test. **** = p value<0.0001. **B-C.** Spheroid growth of Ctrl and DKO A549 cells in ultra-low attachment plates was performed for 13 days and treated with 2 μM ADAM10 inhibitor (GI) or 2μM ADAM17/ADAM10 inhibitor (GW) when indicated. Cell viability quantified using CellTiter Glo was normalised to Ctrl. At least three biological replicates were performed per condition, error bars represent standard deviations and statistical tests were performed using two-tailed student t-test. ns = p value>0.05, **** = p value<0.0001. Representative spheroids of B are shown as insets, scale bar = 0.2 mm. **D.** Tumour volume of Ctrl and iRhom1/2 DKO A549 xenografts assessed twice weekly, starting 7 days post injection of 10^6^ cells in immunodeficient NSG mice (n=6 mice per cell line). Error bars represent standard errors of the mean and statistical tests were performed using two-tailed student t-test. ** = p value<0.01, *** = p value<0.001. Insets show representative tumours, scale bar = 5 mm.

We next assayed the requirement for iRhoms in the growth of A549 spheroids, 3-dimensional models of solid tumours (52, 53). Supporting the significance of the standard 2D cell culture result (Fig. S4B), loss of iRhom1 and iRhom2 also significantly inhibited spheroid growth (Fig. 4B). Consistent with the implication that iRhom-induced release of ERBB ligands contributes to spheroid growth, inhibition of ADAM17 but not ADAM10 also inhibited growth (Fig. 4C, S4C).

These results prompted us to ask whether iRhoms also participate in tumorigenesis *in vivo*, using a xenograft model in which A549 cells are injected into immunodeficient mice. This xenograft model allows preclinical evaluation of the role of candidate target genes in tumour formation and maintenance (54). We established xenografts of A549 parental cells and A549-DKO cells, and found that loss of iRhoms had a profound effect, preventing all detectable tumour growth (Fig. 4D). We conclude that in three models of lung cancer, A549 cells in 2D cell culture, 3D spheroid growth, and tumour xenografts, iRhoms are required for oncogenic signalling and tumour growth.

### iRhom2 phosphorylation regulates ADAM17-dependent release of ERBB ligand and tumour spheroid growth in lung cancer cells

Having established that iRhoms are required in lung tumorigenesis models, we addressed the molecular mechanism that underlies the pro-tumorigenic function of iRhom2 in A549 cells, using our earlier work in HEK293T cells as a guide. First, we made a phosphomutant version of iRhom2 in which the important phosphorylation sites were mutated to alanine (iRhom2^pMUT^); these changes significantly inhibited the release from A549 cells of endogenous amphiregulin (Fig. 5A, B). Second, ERK1/2 kinases drive this mechanism, as the inhibitor U1026 blocked this release (Fig. S5A). Third, phosphorylation of iRhom2 is required for 14-3-3 binding in A549 cells (Fig. 5C), indicating that the phosphorylated iRhom2/14-3-3/ADAM17 activation pathway controls shedding of the ERBB ligands in these lung cancer cells. Together with our results in HEK293T cells, these results support the conclusion that oncogenic KRAS drives ERBB signalling by inducing iRhom2 phosphorylation. As ERBB signalling has recently been shown to contribute to lung tumorigenesis, including in A549 xenograft tumours (29, 30), our results highlight the pro-tumorigenic role of iRhom2-dependent shedding of ERBB ligands in lung cancer cells. We directly tested this conclusion using the spheroid assay, which showed that spheroid growth of A549-DKO cells was significantly reduced in iRhom2^pMUT^ expressing cells, compared to cells expressing iRhom2^WT^ (Fig. 5D).

**Figure 5.**
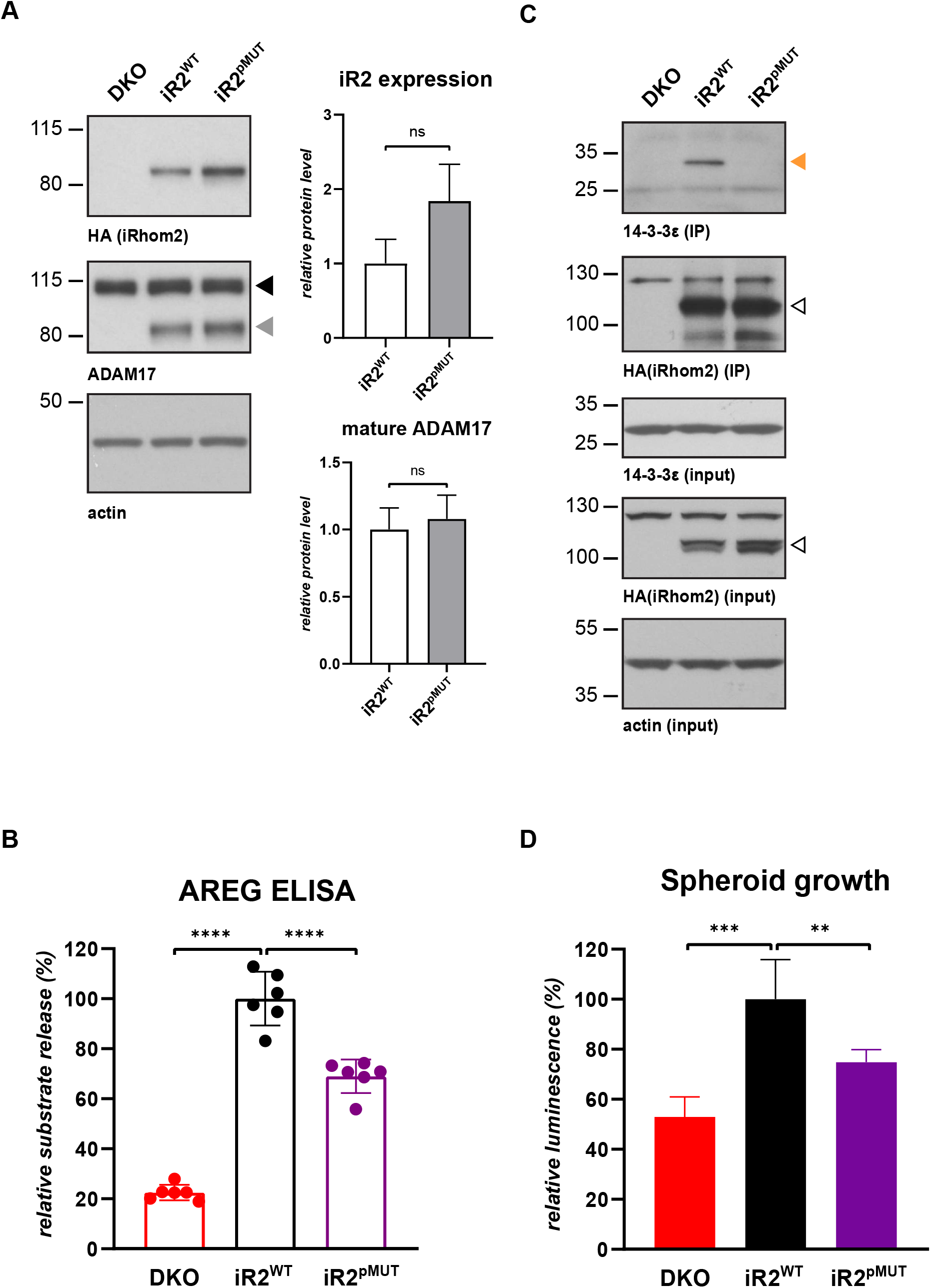
iRhom2 phosphorylation regulates ADAM17-dependent release of ERBB ligand and tumour spheroid growth in lung cancer cells. **A.** iRhom1/2 DKO A549 cells reconstituted with HA-tagged iRhom2^WT^ or phosphomutant iRhom2 (iRhom2^pMUT^) were immunoblotted for HA, ADAM17 and beta-actin. Grey and black arrowheads indicate mature and immature ADAM17 respectively. iRhom2 and mature ADAM17 levels from five biological replicates were quantified relative to beta-actin level using ImageJ. **B.** Release of endogenous amphiregulin (AREG) from DKO A549 parental cells, or those stably expressing iRhom2^WT^ or iRhom2^pMUT^ was measured in six biological replicates by ELISA after overnight collection and normalised as described previously. Error bars represent standard deviations and statistical tests were performed using two-tailed student t-test. **** = p value<0.0001. **C.** HA immunoprecipitates and lysates from A549 DKO cells stably expressing HA-tagged iRhom2^WT^ or iRhom2^pMUT^, immunoblotted for 14-3-3ε, HA and actin. Orange and open arrowheads indicate 14-3-3ε and HA-tagged iRhom2 constructs respectively. This experiment was performed in biological triplicates. **D.** Spheroid growth of DKO A549 parental cells, or those stably expressing iRhom2^WT^ or iRhom2^pMUT^ was measured after 14 days in five biological replicates by CellTiter Glo and normalised as described previously. Error bars represent standard deviations and statistical tests were performed using two-tailed student t-test. ** = p value<0.01, *** = p value<0.001.

### Cancer-associated mutations in iRhom2 increase RAS activity and drive a positive feedback loop in lung cancer cells

In HEK293T cells the cancer causing tylotic iRhom2 mutant D188N enhanced amphiregulin release by oncogenic KRAS mutations (Fig. 3). The same experiment in A549 cells confirmed this result in the lung cancer cell line: compared to iRhom2^WT^, expressing iRhom2^D188N^ in A549-DKO cells caused a more than two-fold increase in the release of endogenous amphiregulin in the presence of oncogenic KRAS (Fig. 6A, B), indicating that tylotic mutation sensitises iRhom2 to oncogenic signalling. Strikingly, iRhom2^D188N^ also further increased spheroid growth compared to iRhom2^WT^ (Fig. 6C), demonstrating that even in transformed A549 cells, the elevated release of ERBB ligand caused by the tylotic iRhom2 mutation was sufficient to further promote tumour-like growth. To emphasise the implication of this result, it demonstrates that a single point mutation in the N-terminus of iRhom2 is sufficient to increase the tumorigenic growth of lung cancer cells.

**Figure 6.**
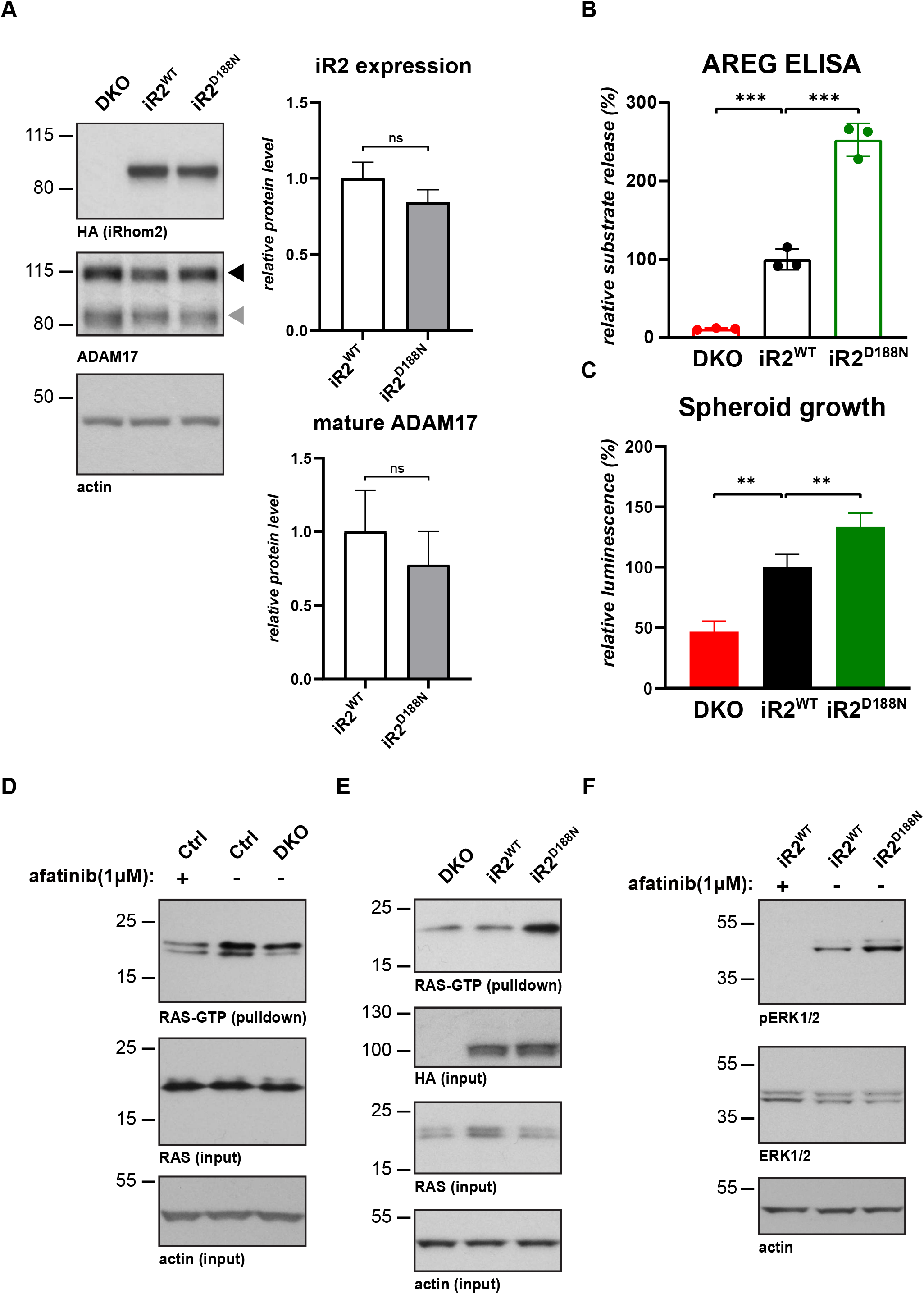
Cancer-associated mutations in iRhom2 increase RAS activity and drive a positive feedback loop in lung cancer cells. **A.** iRhom1/2 DKO A549 cells reconstituted with HA-tagged TOC iRhom2^D188N^ or iRhom2^WT^ were immunoblotted for HA, ADAM17 and beta-actin. Grey and black arrowheads indicate mature and immature ADAM17 respectively. iRhom2 and mature ADAM17 levels from three biological replicates were quantified relative to beta-actin level using ImageJ. **B.** Release of endogenous amphiregulin (AREG) from DKO A549 parental cells, or those stably expressing iRhom2^WT^ or iRhom2^D188N^, was measured in three biological replicates by ELISA after overnight medium collection and normalised as described previously. Error bars represent standard deviations and statistical tests were performed using two-tailed student t-test. *** = p value<0.001. **C.** Spheroid growth of DKO A549 parental cells, or those stably expressing iRhom2^WT^ or iRhom2^D188N^, was measured after 14 days in five biological replicates by CellTiter Glo and normalised as described previously. Error bars represent standard deviations and statistical tests were performed using two-tailed student t-test. ** = p value<0.01. **D-E.** Active RAS was assayed in Ctrl, iRhom1/2 DKO parental A549 cells, or DKO cells stably expressing HA-tagged iRhom2^WT^ or TOC iRhom2^D188N^ using RAS-GTP pulldown. Cells were treated with 1 μM the pan-ERBB inhibitor afatinib for 20 hours when indicated, and immunoblotted for RAS, HA or beta-actin. The experiments were performed in biological triplicates. **F.** Conditioned medium from iRhom1/2 DKO A549 cells stably expressing iRhom2^WT^ or TOC iRhom2^D188N^ was used to stimulate the ERBB1/EGFR reporter cell line A431 treated with 1 μM afatinib when indicated. Following stimulation, A431 cells were immunoblotted for ERK1/2, phosphorylated ERK1/2 (pERK1/2) and beta-actin. The experiment was performed in biological triplicates.

Our observation that iRhoms, and in particular tylotic iRhom2^D188N^, induce ERBB signalling, suggests the existence of a tumorigenic positive feedback loop: oncogenic KRAS, signalling through iRhom2, ADAM17 and amphiregulin, promotes ERBB activity and ultimately further KRAS activity. This possibility builds on recent results that show that oncogenic KRAS mutations are not fully constitutive: using an allele-specific inhibitor it was shown that the activity of KRAS mutant is modulated by upstream ERBB signalling (55, 56). To test this hypothesis, we assayed the activity of oncogenic KRAS in A549 cells by using the RAS-binding domain of Raf1 to pulldown active RAS^GTP^. Compared to parental cells, RAS^GTP^ was as reduced by the absence of iRhoms in A549-DKO as upon the treatment with the pan-ERBB inhibitor afatinib (Fig. 6D, S6A), thus suggesting that iRhoms are required to maintain RAS activity by activating ERBB signalling. As tylotic iRhom2^D188N^ triggers a strong increase in RAS^GTP^ (Fig. 6E, S6B), it further establishes the central role of iRhoms in controlling RAS activity. To definitively conclude whether this feedback loop acts through iRhom2-dependent shedding in the extracellular medium, we assessed the effect of conditioned medium from A549 cells on the ERBB1/EGFR reporter cell line A431. Conditioned medium from tylotic iRhom2^D188N^ caused elevated activated ERK1/2 compared to iRhom2^WT^ (Fig. 6F, S6C). We confirmed that iRhom2-driven activation of the Raf/MEK/ERK pathway depends on ERBB signalling by using afatinib (Fig. 6F, S6C). Together, these results support that iRhom-dependent shedding of ERBB ligands in the extracellular medium drives a positive feedback loop to maintain the activity of oncogenic KRAS in lung cancer cells.

## Discussion

We have discovered that, by regulating ADAM17-dependent release of ERBB ligands, iRhoms are required for KRAS-driven tumorigenesis. Oncogenic mutants of KRAS induce ERK1/2-dependent phosphorylation of the cytoplasmic domain of iRhom2, triggering the recruitment of the phospho-binding proteins 14-3-3, which in turn activate ADAM17 to shed ERBB ligands from the plasma membrane (Fig. 7) The relevance of this mechanism to human disease is demonstrated by our discovery that mutations in the cytoplasmic domain of iRhom2, known to be causative of the human cancer syndrome TOC, are sufficient to amplify this signalling pathway. The significance of iRhom2 to cancer pathogenesis is further reinforced by the result that loss of iRhom activity from A549 lung cancer cells completely blocks their ability to form tumours in a xenograft model.

**Figure 7.**
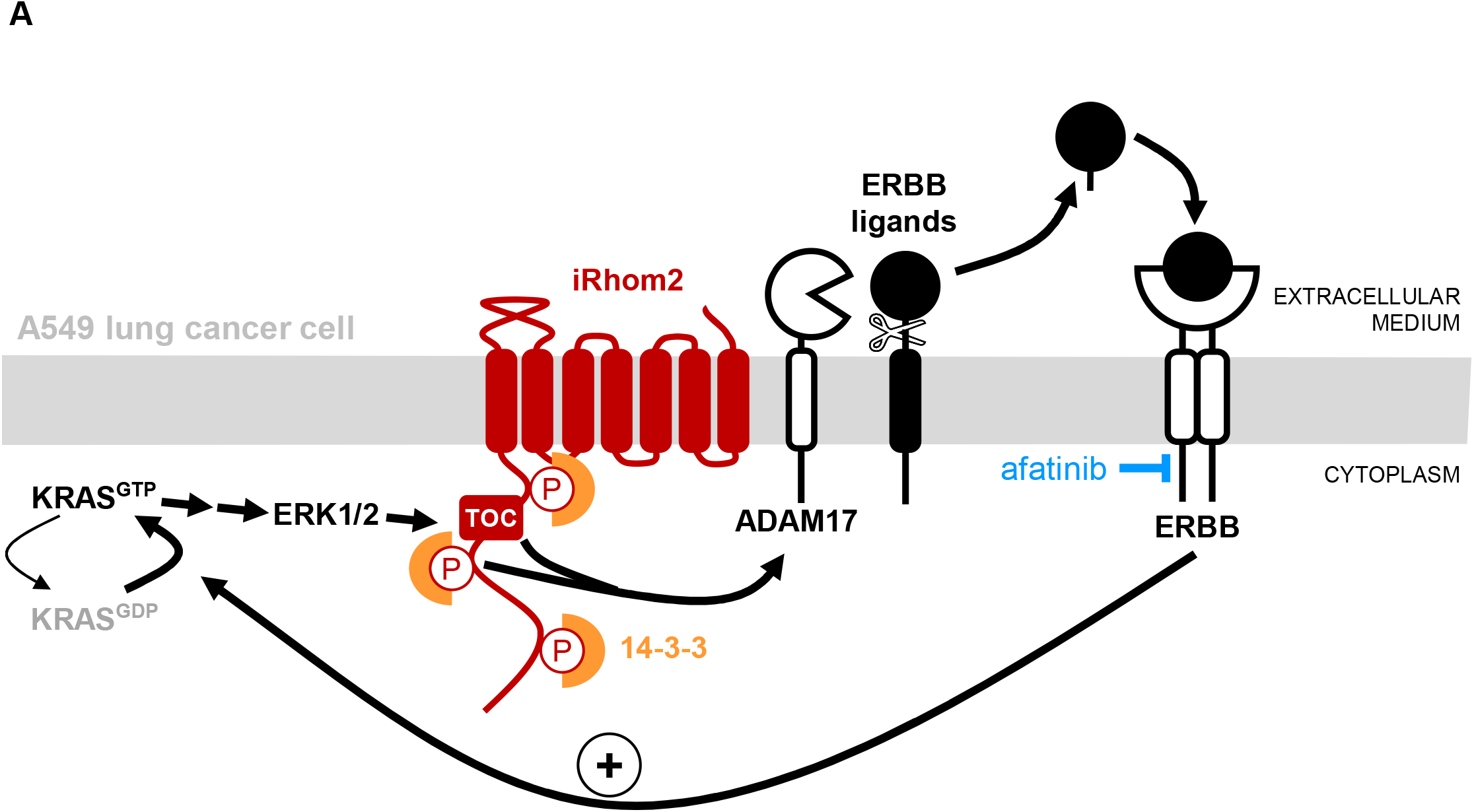
iRhom activity drives an ERBB-dependent feedback loop on oncogenic KRAS. **A.** Activation of ERK1/2 by oncogenic KRAS^GTP^ triggers the phosphorylation of iRhom2 and subsequent recruitment of the phospho-binding proteins 14-3-3. Together with mutations responsible for tylosis with oesophageal cancer (TOC), this induces the ADAM17-dependent release of ERBB ligands into the extracellular medium. Upon binding their ligands, ERBBs maintain KRAS in the active GTP-bound state, thus enabling a positive feedback loop for KRAS oncogenesis. This feedback loop can be inhibited by blocking ERBB signalling with afatinib.

As well as identifying iRhom2 as an essential player in KRAS-induced tumorigenesis, these results reveal the existence of a previously unidentified positive feedback loop that maintains RAS activity in lung cancer cells. In agreement with biochemical evidence proving that, contrary to prior belief, activated KRAS mutations are ‘hyperexcitable’ rather than constitutively locked in an active state (55, 56), two recent studies have shown that oncogenic KRAS relies on upstream ERBB signalling to remain active, and thus to drive lung tumorigenesis (29, 30). Our data establish that the cytoplasmic domain of iRhom2 is crucial in this mechanism: by being both downstream of oncogenic KRAS, and sufficient to increase ERBB-dependent RAS activation, the cytoplasmic domain of iRhom2 represents a central component of this newly uncovered positive feedback loop (Fig. 7). The existence of this feedback mechanism presupposes a sufficient pool of immature, plasma membrane-bound ERBB ligands that can be released in response to elevated iRhom2/ADAM17 activity to reinforce oncogenic KRAS activity. This requirement is supported by the recent observation that the expression of amphiregulin (and other ERBB ligands) is indeed elevated in KRAS-induced lung tumours (29). Overall, our results strengthen a now compelling body of evidence that overturns the earlier belief that oncogenic KRAS mutations are fully constitutive: instead, it is clear that KRAS-driven tumours are driven by signalling input to the activated KRAS oncoprotein. This opens a new potential strategy for therapeutic intervention.

Our results demonstrate that oncogenic and inflammatory signalling pathways share a conserved mechanism for the activation of the iRhom2/ADAM17 complex (this study and (13, 14)). One molecular aspect of the activation of ADAM17 by iRhom2 that we previously reported was that phosphorylation and 14-3-3 binding to iRhom2 causes some kind of conformational change in the complex between the two proteins, detected by weaker binding between them (14). This partial uncoupling also occurred during KRAS-induced shedding (Fig. 2C), thus further demonstrating the conserved activation of the iRhom2/ADAM17 complex. Finally, the growing number of functional signalling complexes in which iRhom2 participates – iRhom2 and ADAM17 (13, 14, 57, 58), iRhom2 and KRAS (Fig. 2C, (59)), and iRhom2 and the previously described binding partner FRMD8 (60, 61) – strengthen the incentives to adopt mechanistic and structural approaches to understanding how iRhom2 controls ADAM17 signalling.

In work that complements these results, Saad et al. reported that phosphorylation of ADAM17 is also important in KRAS-induced lung tumorigenesis (31). They demonstrated that oncogenic KRAS induces the phosphorylation of ADAM17, leading to the shedding of soluble IL-6R and an increase of ERK1/2 activation. Together with our work demonstrating a pathway dependent on phosphorylation of iRhom2 that leads to shedding of ERBB ligands, this establishes the wider significance of MAPK-induced shedding by ADAM17 as a mediator of oncogenic KRAS signalling. It will be interesting to explore the differences and possible crosstalk between the systems that lead, on one hand to shedding of soluble IL-6R triggered by phosphorylated ADAM17, and on the other, to ERBB ligand shedding induced by phosphorylated iRhom2.

Oncogenic KRAS is a driver of multiple cancers in addition to lung adenocarcinoma, so our work raises the question of whether iRhom2 also has a role in these other cancers. Pancreatic adenocarcinoma, the seventh leading cause of cancer-related deaths worldwide, is considered the most KRAS-addicted cancer (62–64). Strikingly, ADAM17 and ERBB1/EGFR are both required to maintain high RAS activity in a *Kras^G12D^* mouse model of pancreatic ductal carcinoma (27, 28). In the light of the results we report here, it will be interesting to investigate whether iRhom2 plays a similar role in supporting a positive feedback loop in this particularly aggressive oncogenic context. In support of this possibility, the cytoplasmic domain of iRhom2 has been found to be phosphorylated in the presence of KRAS^G12D^ in pancreatic cancer cells (65). Another case where there is now a strong incentive to explore the possible involvement of iRhom2 is colorectal cancer, the second leading cause of cancer-related deaths worldwide (62, 66), which can also be driven by oncogenic KRAS mutations (67). Using patient-derived organoids and xenografts it has recently been demonstrated that ERBB signalling promotes tumorigenesis by maintaining ERK activity in colorectal tumours (68). Although ADAM17 has been shown to be required for colorectal tumour growth (10), the possible contribution of iRhom2 phosphorylation in colorectal tumorigenesis is currently unexplored.

In summary, we have shown that by driving ADAM17-dependent ERBB signalling, iRhoms are essential components in KRAS-driven tumorigenesis. On a mechanistic level, we report the existence of a KRAS-iRhom2-ERBB positive feedback loop that maintains oncogenic KRAS activity and may explain the potency of KRAS-induced cancers. Finally, by establishing the role of iRhom2 in oncogenic activation of ADAM17, our results provide new routes to explore future therapeutic opportunities.

## Material and Methods

### Molecular cloning

iRhom2, KRAS4A, KRAS4B SRC, BRAF and AKT1 constructs were amplified by PCR from *iRhom2* cDNA (60), *KRAS4A* cDNA (69), *KRAS4B* cDNA (kind gift from Julian Downward (Francis Crick Institute, London)), *SRC* cDNA (antibodies-online), *BRAF* cDNA (antibodies-online) and *AKT1* cDNA (antibodies-online). They were mutated using QuikChange Multi Site-Directed Mutagenesis Kit (Agilent Technologies, 200515) and subcloned using In-Fusion HD Cloning Kit (Takara Bio, 639649) according to the manufacturer’s instructions. For all constructs, single colonies were picked and extracted DNA was verified by Sanger sequencing (Source Bioscience, Oxford, UK).

### List of plasmids

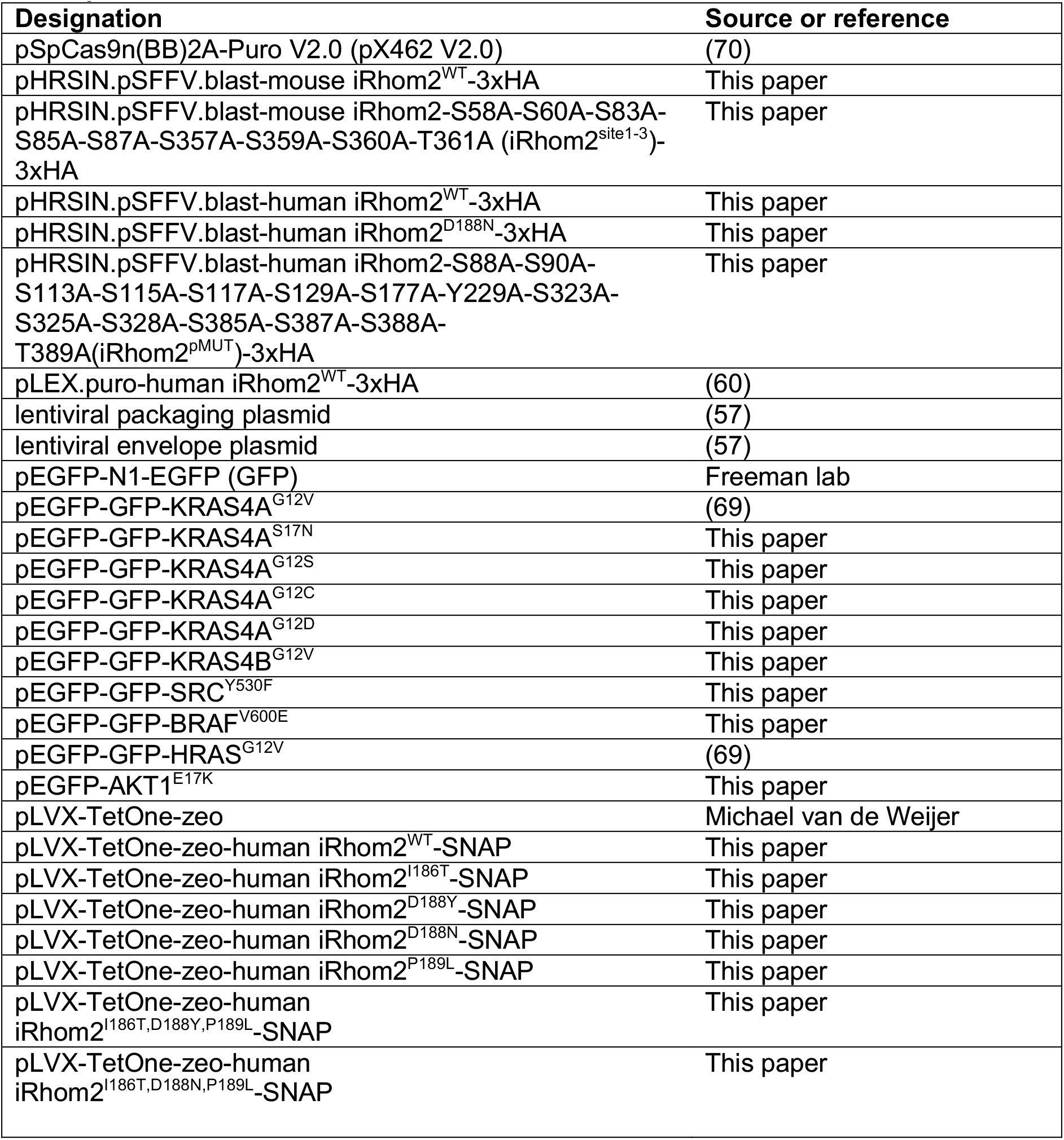

### Cell culture and DNA Transfection

Human embryonic kidney (HEK) 293T cells and human non-small-cell lung cancer (NSCLC) A549 cells were cultured in DMEM (Sigma-Aldrich) supplemented with 10% fetal bovine serum (FBS) (Sigma-Aldrich) and 2 mM L-Glutamine (Gibco) at 37°C with 5% CO2. Human carcinoma A431 cells were cultured in EMEM (Lonza) supplemented with 10% FBS and 2 mM L-Glutamine (Gibco). FuGENE HD (Promega) was used for transient DNA transfection in HEK293T cells, with a ratio of 1 μg DNA and 4 μl transfection reagent diluted in OptiMEM (Gibco). Lipofectamine 2000 (Thermo Fisher Scientific) was used for transient DNA transfection of A549 cells, with a ratio of 0.3 μg DNA and 1 μl transfection reagent. HEK DKO stably expressing pLVX-TetOne-zeo constructs were stimulated with 100 ng/ml doxycycline (MP Biomedicals, 195044).

### CRISPR/Cas9 genome editing in A549 and HEK293T cells

CRISPR/Cas9-mediated single knockout of human *RHBDF1/iRhom1* or *RHBDF2/iRhom2* in HEK293T was performed as described before (60). In brief, the plasmids co-expressing Cas9 nickase (Cas9n) and the gRNA targeting *RHBDF1* or *RHBDF2* were transfected into HEK293T cells. Upon puromycin selection and isolation of single colonies, the loss of *RHBDF1* or *RHBDF2* was analysed by PCR.

For CRISPR/Cas9-mediated double knockout of human *RHBDF1/iRhom1* and *RHBDF2/iRhom2* in A549 cells, 4 μg of plasmids co-expressing Cas9n and the gRNA were transfected using the Neon Transfection System (Invitrogen) according to the manufacturers’ instructions. The following electroporation settings were used: 1,230 volts, 30 seconds pulse width, 2 pulses number and 8 x 10^6^ cells/ml. Antibiotic selection was performed using 0.5 μg/ml puromycin for 48 hrs, before selecting single colonies to establish clonal cell lines, and analysing loss of *RHBDF1* and *RHBDF2* by PCR.

### List of primers

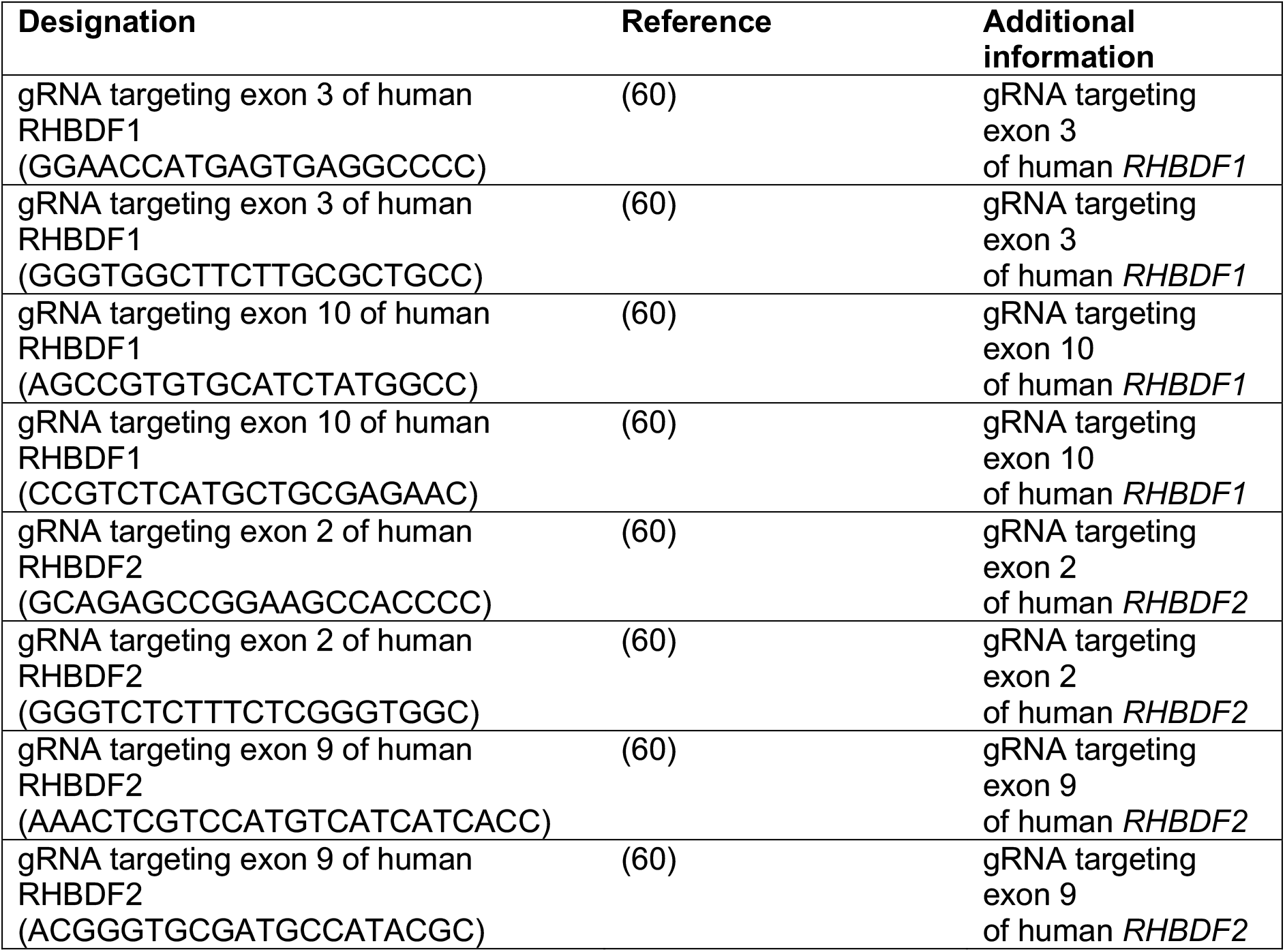

### Lentiviral transduction of cell lines

A549 or HEK293T DKO cells stably expressing iRhom2 constructs were generated by lentiviral transduction using the pLVX-TetOne or pHRSIN constructs as previously described (57). Cells were selected by adding 50 μg/ml zeocin or 10 μg/ml Blasticidin S HCl.

### List of cell lines

| Designation | Source or reference | Additional information |
| --- | --- | --- |
| HEK293T cells | Freeman lab |  |
| HEK293T iRhom1/iRhom2 double-knockout (DKO) | (60) | CRISPR/Cas9-mediated KO cell line |
| HEK293T iRhom1/iRhom2 DKO + iRhom2<sup>WT</sup> | (60) | HEK293T DKO cells transduced with pLEX.puro-human iRhom2<sup>WT</sup>-3xHA |
| HEK293T iRhom1 knockout | This paper | CRISPR/Cas9-mediated KO cell line |
| HEK293T iRhom2 knockout | This paper | CRISPR/Cas9-mediated KO cell line |
| HEK293T iRhom1/iRhom2 DKO + iRhom2<sup>WT</sup> | This paper | HEK293T DKO cells transduced with pHRSIN.pSFFV.blast-mouse iRhom2<sup>WT</sup>-3xHA, used as control for HEK-DKO-iRhom2<sup>site1-3</sup> |
| HEK293T iRhom1/iRhom2 DKO + iRhom2<sup>site1-3</sup> | This paper | HEK293T DKO cells transduced with pHRSIN.pSFFV.blast-mouse iRhom2<sup>site1-3</sup>-3xHA |
| HEK293T iRhom1/iRhom2 DKO + iRhom2<sup>WT</sup> (inducible) | This paper | HEK293T DKO cells transduced with pLVX-TetOne-zeo-human iRhom2<sup>WT</sup>-SNAP, used as control for HEK DKO expressing iRhom2 TOC constructs |
| HEK293T iRhom1/iRhom2 DKO + iRhom2<sup>I186T</sup> (inducible) | This paper | HEK293T DKO cells transduced with pLVX-TetOne-zeo-human iRhom2<sup>I186T</sup>-SNAP |
| HEK293T iRhom1/iRhom2 DKO + iRhom2<sup>D188Y</sup> (inducible) | This paper | HEK293T DKO cells transduced with pLVX-TetOne-zeo-human iRhom2<sup>D188Y</sup>-SNAP |
| HEK293T iRhom1/iRhom2 DKO + iRhom2<sup>D188N</sup> (inducible) | This paper | HEK293T DKO cells transduced with pLVX-TetOne-zeo-human iRhom2<sup>D188N</sup>-SNAP |
| HEK293T iRhom1/iRhom2 DKO + iRhom2<sup>P189L</sup> (inducible) | This paper | HEK293T DKO cells transduced with pLVX-TetOne-zeo-human iRhom2<sup>P189L</sup>-SNAP |
| HEK293T iRhom1/iRhom2 DKO + iRhom2<sup>I186T,D188Y,P189L</sup> (inducible) | This paper | HEK293T DKO cells transduced with pLVX-TetOne-zeo-human iRhom2<sup>I186T,D188Y,P189L</sup>-SNAP |
| HEK293T iRhom1/iRhom2 DKO + iRhom2<sup>I186T,D188N,P189L</sup> (inducible) | This paper | HEK293T DKO cells transduced with pLVX-TetOne-zeo-human iRhom2<sup>I186T,D188N,P189L</sup>-SNAP |
| A549 cells | (71) |  |
| A549 iRhom1/iRhom2 DKO | This paper | CRISPR/Cas9-mediated KO cell line |
| A549 iRhom1/iRhom2 DKO + iRhom2<sup>WT</sup> | This paper | A549 DKO cells transduced at high multiplicity of infection (MOI) with pHRSIN.pSFFV.blast-human iRhom2<sup>WT</sup>-3xHA, used as control for A549-DKO-iRhom2<sup>D188N</sup> |
| A549 iRhom1/iRhom2 DKO + iRhom2<sup>D188N</sup> | This paper | A549 DKO cells transduced with pHRSIN.pSFFV.blast-human iRhom2<sup>D188N</sup>-3xHA |
| A549 iRhom1/iRhom2 DKO + iRhom2<sup>WT</sup> | This paper | A549 DKO cells transduced at low MOI with pHRSIN.pSFFV.blast-human iRhom2<sup>WT</sup>-3xHA, used as control for A549-DKO-iRhom2<sup>pMUT</sup> |
| A549 iRhom1/iRhom2 DKO + iRhom2<sup>pMUT</sup> | This paper | A549 DKO cells transduced with pHRSIN.pSFFV.blast-human iRhom2<sup>pMUT</sup>-3xHA |
| A431 cells | Freeman lab |  |

### Co-immunoprecipitation

Cells were washed three times with ice-cold PBS before lysis in Triton X-100 lysis buffer (1% Triton X-100, 150 mM NaCl, 50 mM Tris-HCl (pH 7.5)) supplemented with EDTA-free complete protease inhibitor mix (Roche, 11873580001) and 10 mM 1,10-phenanthroline (Sigma-Aldrich, 131377–5G). Pre-washed anti-HA magnetic beads (Thermo Scientific, 88837) were added to the lysates cleared from cell debris by centrifugation at 15,000 rpm at 4°C for 15 min and incubated for at least 2 hr on a rotor at 4°C. Beads were washed five times with Triton X-100 lysis buffer and eluted with a 10-minute incubation at 65°C in 2x SDS sample buffer (0.25 M TrisHCl pH6.8, 10% SDS, 50% glycerol, 0.02% bromophenol blue) supplemented with 200 mM DTT.

### Concanavalin A enrichment

Cell lysates were incubated with 30 μl concanavalin A sepharose (Sigma-Aldrich, C9017-25ML) at 4°C for 2 hr on a rotor. Beads were pelleted at 4000 rpm for 2 min at 4°C and washed five times with Triton X-100 lysis buffer. Glycoroteins were eluted with 2x LDS buffer (Invitrogen) supplemented with 25% sucrose and 50 mM DTT for 10 min at 65°C.

### RAS-GTP pulldown

To detect active RAS in A549 cells, RAS-GTP pulldown was performed according to the manufacturers’ instructions using the Active Ras Detection Kit (Cell Signaling Technology, #8821). In brief, one confluent 10 cm dish of cells was rinsed with ice-cold PBS and lysed in 0.5 ml ice-cold lysis buffer supplemented with 1 mM PMSF. Cell lysates were cleared by centrifugation and protein concentration was determined by Bradford assay. Cleared lysates were added to the pre-washed spin cup which contains 100 μl of the 50% resin slurry and 80 μg of GST-Raf1-RBD and incubated at 4°C for 1 hr on a rotor. The resin was washed three times with Wash Buffer and the proteins bound to the resin were eluted with 50 μl of the sample buffer supplemented with 200 mM DTT. Samples were denatured at 95°C for 5 min and were subjected to western blot analysis.

### SDS-PAGE and western blotting

Cell lysates were denatured at 65°C for 10 min in sample buffer supplemented with 100 mM DTT. Samples were run in 4-12% Bis-Tris NuPAGE gradient gels (Invitrogen) and MOPS running buffer (50 mM MOPS, 50 mM Tris, 0.1 % SDS, 1 mM EDTA), or in Novex 8-16% Tris-Glycine Mini Gels with WedgeWell format (Thermo Scientific) and Tris-Glycine running buffer (25 mM Tris,192 mM glycine, 0.1% SDS). Proteins were then transferred to a methanol activated polyvinylidene difluoride (PVDF) membrane (Millipore) in Bis-Tris or Tris-Glycine transfer buffer. 5% milk in PBST (0.1% Tween 20) or TBST (0.05% Tween 20) was used for blocking and antibody incubation, and PBST or TBST was used for washing. The membranes were incubated with secondary antibodies at the room temperature for 1 hr. Blots were quantified using ImageJ.

### List of antibodies

| Name | Source | Catalogue number | Dilution |
| --- | --- | --- | --- |
| anti-ADAM17, rabbit polyclonal | Abcam | ab39162 | 1:2000 |
| anti-beta-actin, mouse monoclonal | Santa Cruz | sc-47778 | 1:2000 |
| anti-14-3-3 epsilon, rabbit polyclonal | CST | 9635 | 1:500 |
| anti-HA-HRP, rat monoclonal (clone 3F10) | Roche | 12013819001 | 1:2000 |
| anti-GFP, chicken polyclonal | Abcam | ab13970 | 1:4000 |
| anti-phosphoERK1/2 (T202/Y204), rabbit monoclonal (clone 197G2) | CST | 4377 | 1:500 |
| anti-ERK1/2, rabbit polyclonal | CST | 9102 | 1:1000 |
| anti-V5, goat polyclonal | Santa Cruz | sc-83849 | 1:2000 |
| anti-AKT, rabbit polyclonal | CST | 9272 | 1:1000 |
| anti-pSRC (Y416), rabbit polyclonal | CST | 2101 | 1:1000 |
| anti-pAKT (S473), rabbit polyclonal | CST | 9271 | 1:1000 |
| anti-calreticulin, rabbit polyclonal | Invitrogen | PA3-900 | 1:2000 |
| anti-transferrin receptor1, mouse monoclonal (clone H68.4) | Invitrogen | 13-6800 | 1:2000 |
| anti-rabbit-HRP, goat polyclonal | CST | 7074 | 1:2500 |
| anti-goat-HRP, mouse monoclonal | Santa Cruz | sc2354 | 1:2500 |
| anti-mouse-HRP, horse polyclonal | CST | 7076 | 1:2500 |
| anti-chicken-HRP, goat polyclonal | Novus Biologicals | NB7303 | 1:10000 |

### AP-shedding assay

HEK293T cell lines were seeded in poly-(L)-lysine (PLL, Sigma-Aldrich) coated 24-well plates in triplicates 24 hours before transfection. 50 ng alkaline phosphatase (AP)-conjugated substrates were transfected with FuGENE HD (Promega, E2312). In KRAS related experiments, 100 ng control plasmids or KRAS plasmids were transfected together with AP-substrates. 24 hrs after transfection, cells were washed twice with PBS and incubated for 18 hr in 300 μl phenolred-free OptiMEM (Gibco, 11058-021) supplemented with 1 μM GW280264X (GW) (Generon, AOB3632-5) or GI254023X (GI) (Sigma, SML0789-5MG) when indicated. For kinase inhibition assay, 300 μl phenolred-free OptiMEM were supplemented with 10 μM U0126 (abcam, ab120241-5mg) and the supernatant was collected after 3 hr. The supernatants were then collected, and cells were lysed in 300 μl Triton X-100 lysis buffer supplemented with EDTA-free protease inhibitor mix (Roche). 100 μl supernatant and 100 μl diluted cell lysates were independently incubated with 100 μl AP substrate p-nitrophenyl phosphate (PNPP) (Thermo Scientific, 37620) at room temperature and the absorbance was measured at 405 nm by a plate reader (SpectraMax M3, Molecular Devices). The percentage of substrate release was calculated by dividing the signal from the supernatant by the total signal (supernatant and cell lysate).

### Spheroid assay

Tumour spheroid were generated as previously described in (72). In brief, 2,500 cells were resuspended in culture medium supplemented with 2.5% growth-factor reduced Matrigel (Scientific Laboratory Supplies #356231) and placed in a 96-well round-bottom ultra-low attachment plate (Corning #7007). Formation of the spheroids was initiated by centrifugation at 1,200 rpm for 4 minutes. After 13 days, tumour spheroids were imaged using a stereoscopic microscope (Leica DFC310 FX), and cell viability was measured using the CellTiter-Glo Cell viability assay (Promega, #G9681) according to the manufacturer’s instructions.

### Cell proliferation assay

To assay cell proliferation in a 2D adherent format, 1,000 cells were seeded in standard 96-well tissue culture plate. After five days, cell viability was measured using the CellTiter-Glo Cell viability assay (Promega, #G9681) according to the manufacturer’s instructions, as previously described in (56).

### A431 ERBB1/EGFR activation assay

1.5×10^6^ A549 or 3×10^6^ A431 cells were seeded in a 10 cm tissue culture dish. After three days, A431 cells were washed once with PBS, and serum-starved in 10ml of OptiMEM supplemented with 1 μM afatinib when indicated, and with 2 μM GW and to prevent growth factor release from A431 cells. The following day, the medium of A431 cells was renewed with OptiMEM supplemented with the same inhibitors, while A549 cells were washed once with PBS before adding 5 ml OptiMEM constituting the conditioned medium. After four hours of collection, A431 cells were incubated with the conditioned medium for three minutes before being placed on ice and lysed as described above.

### ELISA

80,000 A549 cells were seeded in triplicates per well of a 24-well plate. To study the loss of shedding in A549-DKO and A549-DKO-iRhom2^pMUT^, the medium was replaced the following day with 350 μl of full medium and collected after 18 hr of incubation. To determine the increased shedding in A549-DKO-iRhom2^D188N^, a 4-hr collection was performed 48 hr after seeding the cells. Similarly, a 4-hr collection in 350 μl of full medium supplemented with 10 μM U0126 was performed to determine the contribution of ERK1/2. In all cases, the concentration of amphiregulin in the supernatant was determined using the Human Amphiregulin Quantikine ELISA Kit (R&D Systems, DAR00) according to the manufacturers’ instructions. In parallel, the cells were lysed in Triton X-100 lysis buffer and the total protein concentration was measured using the BCA Assay (Life Technologies). The substrate release was determined by normalising the amphiregulin concentration by the total protein concentration.

### Xenograft model

1×10^6^ Ctrl and iRhom1/2 double-knockout (DKO) A549 cells were resuspended in Matrigel:PBS (50:50 v/v) before being subcutaneously injected in one flank of 12 (n=6 mice per cell line) 6-week-old female immunodeficient NOD.*Cg-Prkdc^scid^Il2rg^tm1Wjl^*/SzJ (NSG) mice (Charles River UK Ltd (Margate, Kent)). Xenograft growth was monitored with a calliper twice weekly; tumour volume was determined using the following formula: (length x width^2^)/2. At the end of the experiment, tumours were collected and photographed. Animal experiments were performed under the Home Office Project Licence PPL30/3395 (A.J. Ryan licence holder).

## Acknowledgments

We are indebted to Ervin Fodor (Dunn School, Oxford) for providing A549 cells, and to Sally Cowley (Dunn School, Oxford) for the help in generating the CRISPR/Cas9 knockout A549 cell lines. We thank the whole Freeman and Ryan groups for contributions during the course of the project and for feedback on the manuscript. We are thankful to Miriam Molina-Arcas and Julian Downward (Francis Crick Institute, London) for the exchange of ideas and for providing the plasmid to express *KRAS4B*, and to Michael van de Weijer (Dunn School, Oxford) for providing the pLVX-TetOne-zeo plasmid.

The Freeman group was supported by a Wellcome Trust Investigator Awards (101035/Z/13/Z and 220887/Z/20/Z). This project was also supported by a Cancer Research UK Development Fund (CRUKDF 0920-MF). B.S. was funded by the Medical Research Council scholarship (1643127). F.L. was supported by the China Scholarship Council-University of Oxford Scholarship (201806240008). A.G.G. received funding from H2020 Marie Skłodowska-Curie Actions fellowship (659166), and A.G.G. and M.F. were funded by the Biotechnology and Biological Sciences Research Council Research grant 440 (BB/RO16771/1). S.M.S. and A.J.R. were funded by the Medical Research Council Programm Grant (MC_UU_00001/6).

## Additional information

The authors declare no competing interests.

## Author contributions

B.S., F.L., A.G.G., A.J.R. and M.F. designed research; B.S., F.L. and S.M.S. performed research; B.S., F.L., S.M.S. and A.G.G. contributed new reagents/analytical tools; B.S., F.L. and S.M.S. analysed data; B.S., F.L., S.M.S., A.G.G., A.J.R. and M.F. wrote the paper; A.J.R. and M.F. supervised the project.

---

**Figure S1.**
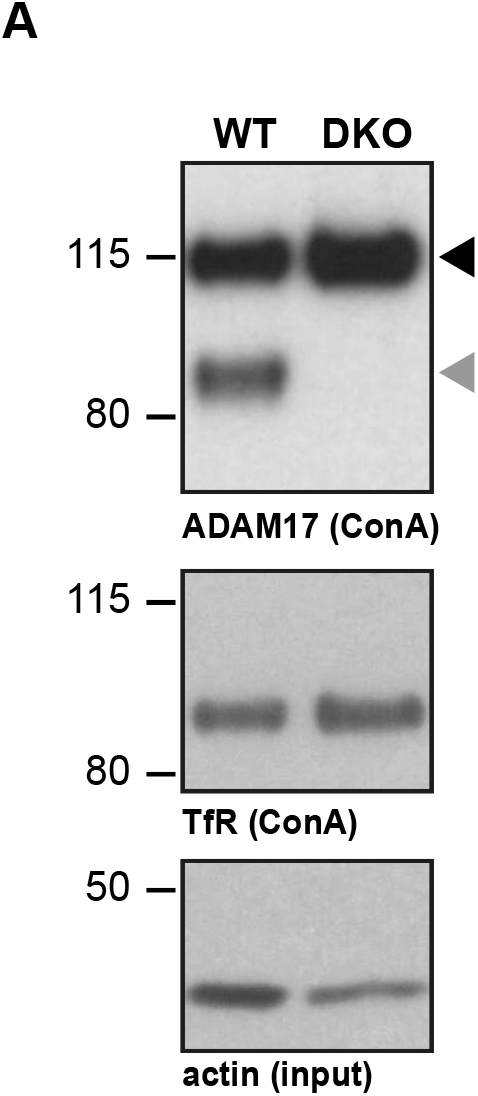
**A.** Concanavalin A (ConA) enrichment of lysates prepared from iRhom1/2 DKO or WT HEK293T and immunoblotted for ADAM17, transferrin receptor (TfR) or beta-actin. The lack of mature ADAM17 (grey arrowhead), compared to immature proADAM17 (black arrowhead), in the absence of iRhoms demonstrates the loss of all iRhom activity. This experiment was repeated three times.

**Figure S2.**
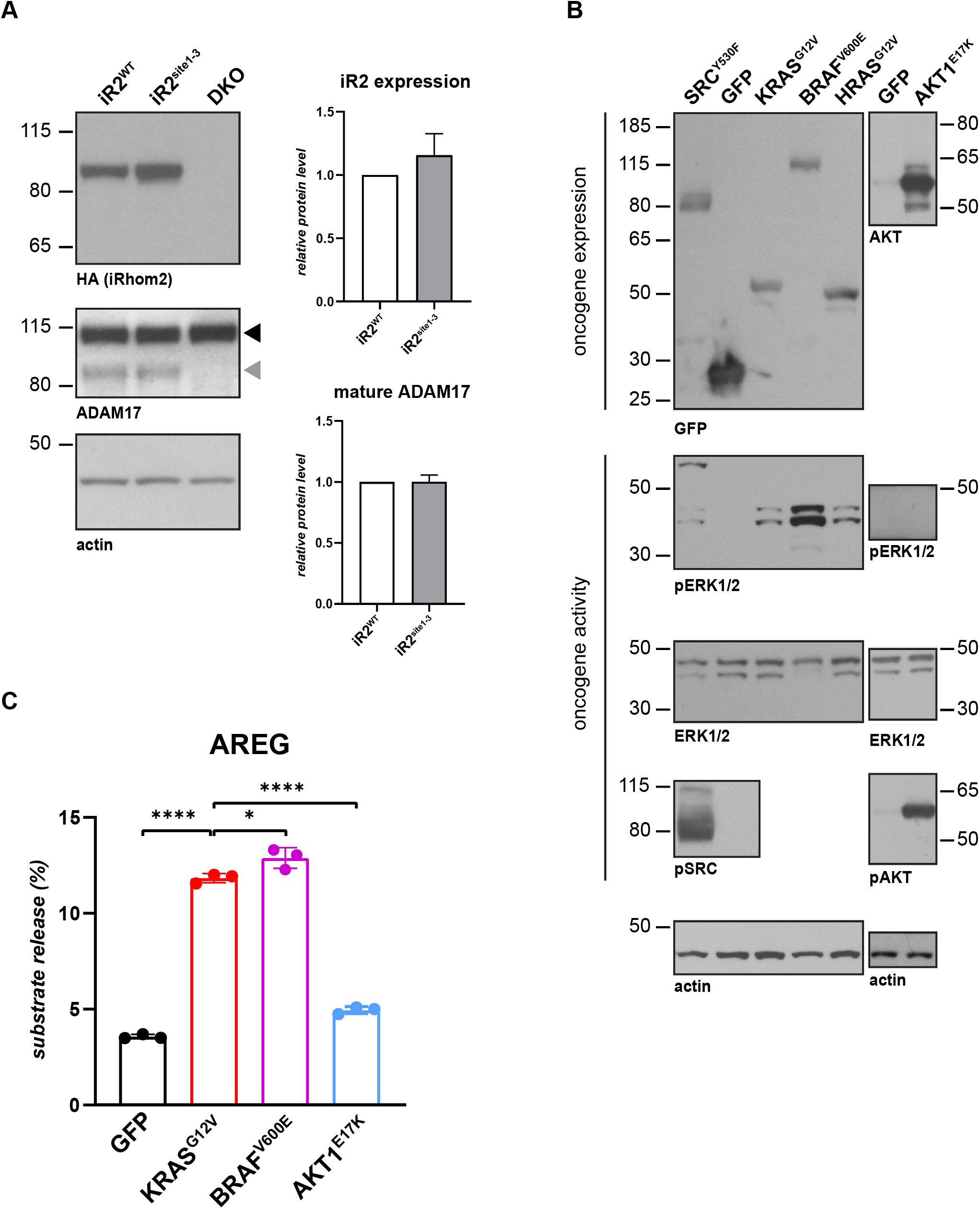
**A.** Lysates from iRhom1/2 DKO HEK293T cells reconstituted with HA-tagged iRhom2^WT^ or iRhom2^site1-3^ were immunoblotted for HA, ADAM17 and beta-actin. Grey and black arrowheads indicate mature and immature ADAM17 respectively. iRhom2 and mature ADAM17 levels from at least three biological replicates were quantified relative to beta-actin level using ImageJ. **B.** HEK293T cells transfected with GFP or GFP-tagged SRC^Y530F^, KRAS^G12V^, BRAF^V600E^, HRAS^G12V^, untagged AKT1^E17K^ were immunoblotted for oncogene expression (GFP and AKT1), induction of phosphorylated ERK1/2 (pERK1/2) or for beta-actin. The level of phosphorylated SRC and AKT were probed as a control of their constitutive activity. The experiment was performed in biological triplicates. **C.** HEK293T cells were transiently co-transfected with AP-tagged AREG and GFP or GFP-tagged KRAS^G12V^, BRAF^V600E^, untagged AKT1^E17K^. Overnight medium collection was performed in three biological replicates. Substrate release is the level of released alkaline phosphatase in the medium divided by the total alkaline phosphatase level. Error bars represent standard deviations and statistical tests were performed using two-tailed student t-test. * = p value<0.05, **** = p value<0.0001.

**Figure S3.**
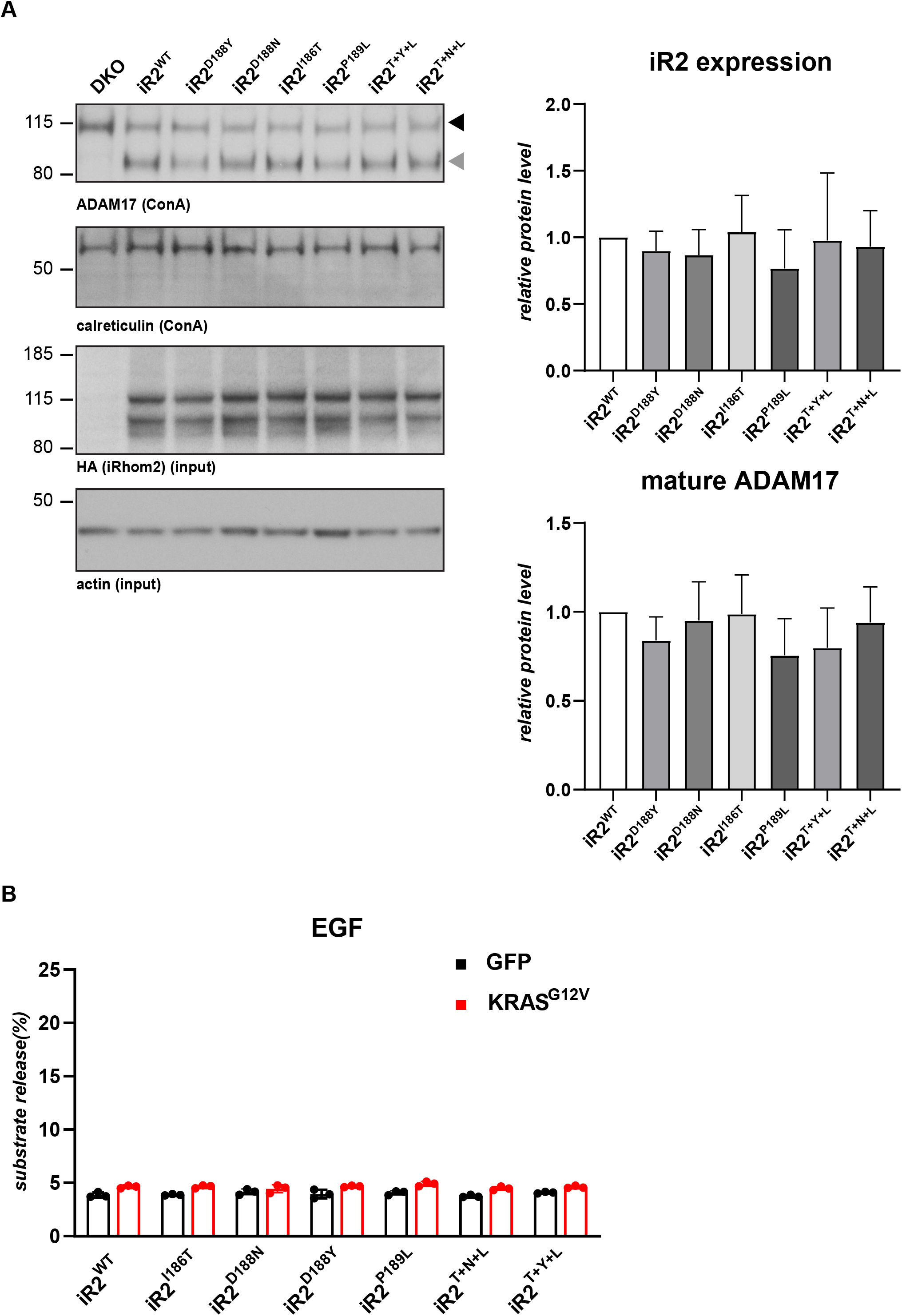
**A.** Concanavalin A (ConA) enrichment of lysates from iRhom1/2 DKO HEK293T cells reconstituted with HA-tagged iRhom2^WT^ or iRhom2 variant harbouring one or a combination of the TOC mutations, followed by immunoblotting for ADAM17 and calreticulin. Black and grey arrowheads indicate immature and mature ADAM17 respectively. Stable expression of HA-tagged iRhom2 variants was detected by HA and beta-actin antibodies. iRhom2 and mature ADAM17 levels from three biological replicates were quantified using ImageJ relative to beta-actin and total ADAM17 (immature and mature) respectively. **B.** iRhom1/2 DKO HEK293T cells reconstituted with iRhom2^WT^ or with an iRhom2 variant harbouring one of the TOC mutations or the three mutations combined: T+Y+L (I186T, D188Y, P189L) or T+N+L (I186T, D188N, P189L) were co-transfected with GFP or GFP-tagged KRAS^G12V^, and alkaline-phosphatase AP-EGF. Overnight collection of medium was performed in biological triplicates.

**Figure S4.**
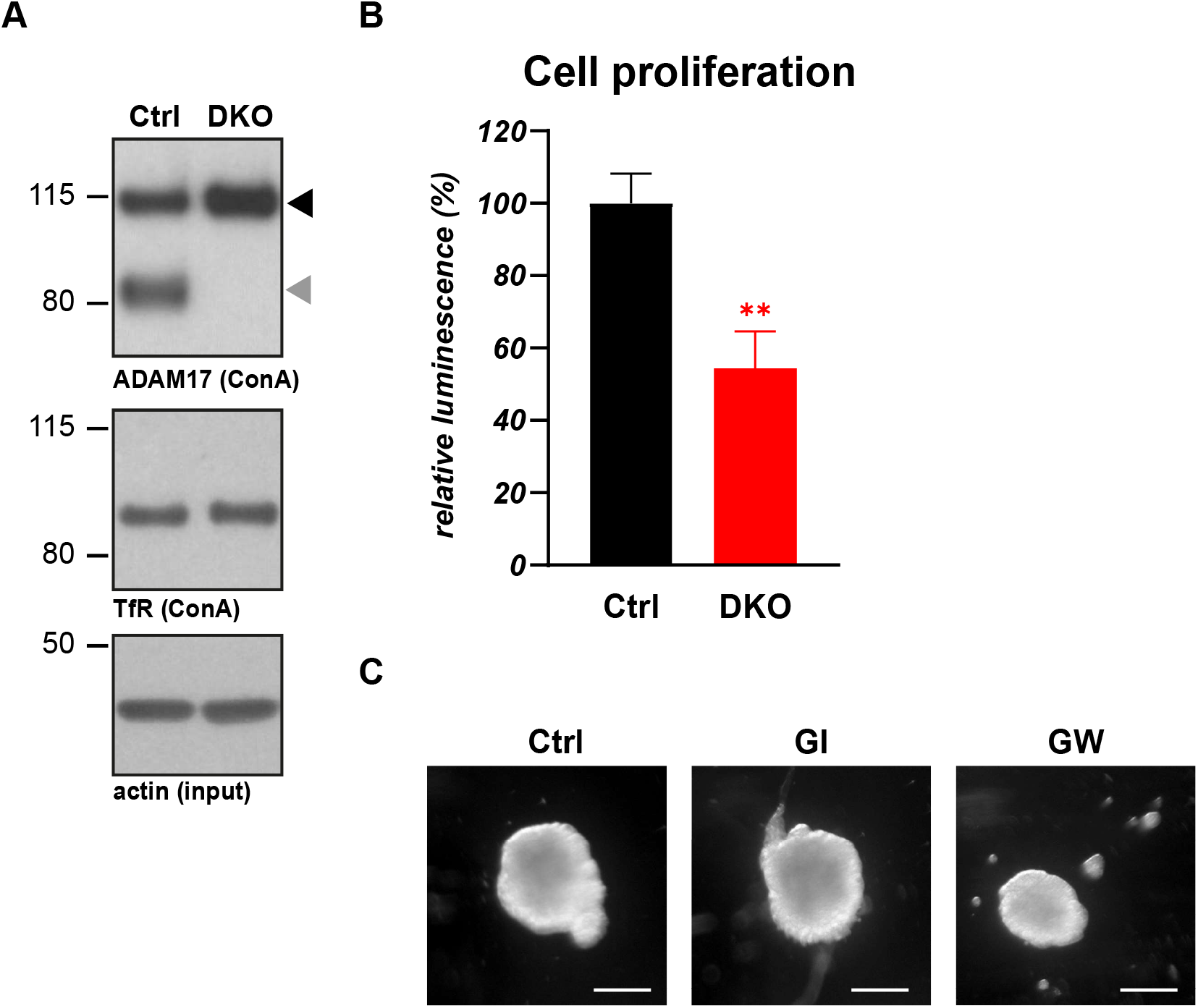
**A.** Concanavalin A (ConA) enrichment of lysates from Ctrl and iRhom1/2 DKO A549 cells, immunoblotted for ADAM17, transferrin receptor (TfR) or beta-actin. The absence of mature ADAM17 (grey arrowhead) compared to immature proADAM17 (black arrowhead) demonstrates the lack of iRhom activity. The experiment was repeated three times. **B.** Cell proliferation of Ctrl and iRhom1/2 DKO A549 cells was measured five days after seeding the cells using CellTiter Glo. The luminescence level was normalised to the level of A549 Ctrl (100%). Three biological replicates were performed per cell line, the error bars represent standard deviations and the statistical tests were performed using two-tailed student t-test. **=p value <0.01. **C.** Representative images of A549 spheroids performed in triplicate are shown after 13 days treatment with 2 μM GI or 2 μM GW when indicated. scale bar = 0.2 mm.

**Figure S5.**
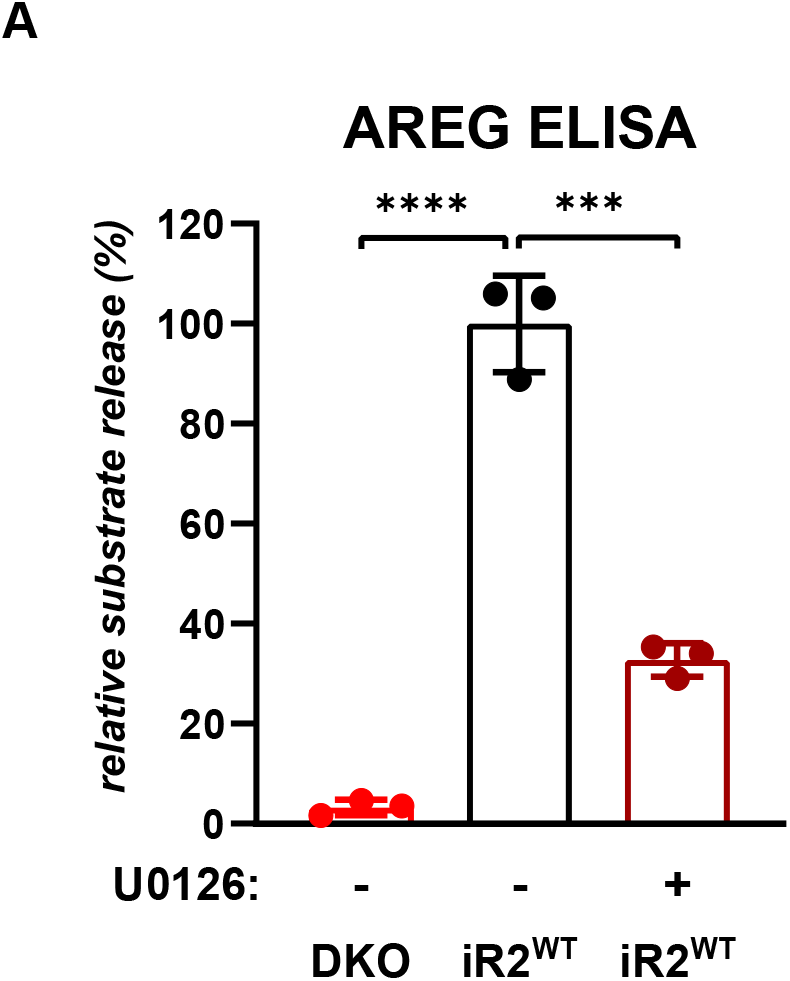
**A.** Release of endogenous AREG from DKO A549 parental cells or those stably expressing iRhom2^WT^, measured by ELISA after four hours of treatment with DMSO or 10 μM U0126. Substrate release was normalised as previously described, error bars represent standard deviations and statistical tests were performed using two-tailed student t-test. *** = p value<0.001, **** = p value<0.0001.

**Figure S6.**
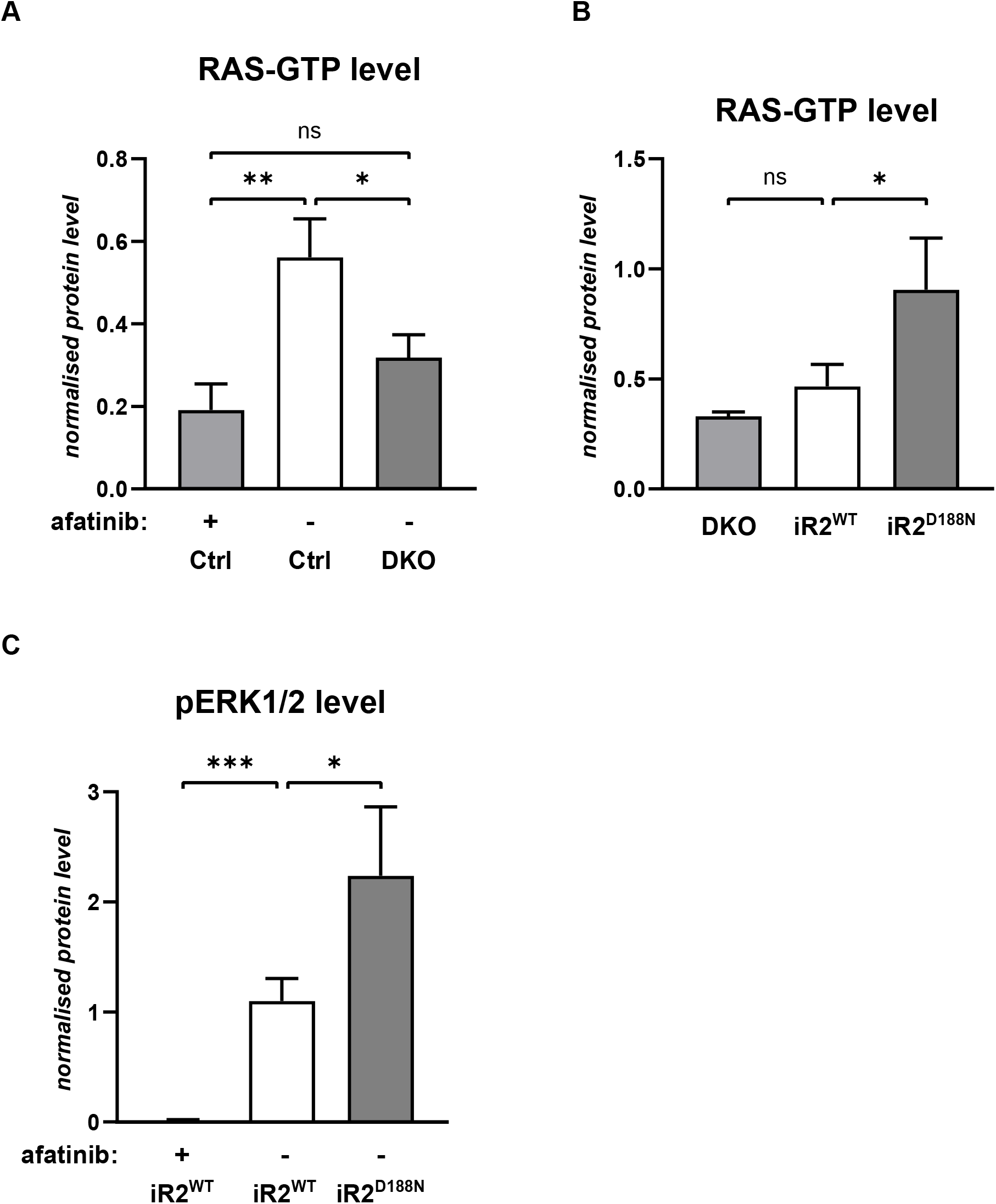
**A-B.** Quantification of active RAS-GTP level from three biological replicates described in Fig. 6D-E was performed using ImageJ. The level of RAS-GTP was normalised to the loading control beta-actin. Error bars represent standard deviations and statistical tests were performed using two-tailed student t-test. ns = p value>0.05, **=p value <0.01. **C.** Quantification of phosphorylated ERK1/2 (pERK1/2) level from three biological replicates described in Fig. 6F was performed using ImageJ. The level of pERK1/2 was normalised on the loading control beta-actin. Error bars represent standard deviations and statistical tests were performed using two-tailed student t-test. *=p value <0.05, *** = p value<0.01.

